# Genome-resolved metagenomics of traditional fermented beverages reveals biosynthetic diversity and informs the rational *in silico* design of probiotic synthetic communities

**DOI:** 10.64898/2026.07.22.740150

**Authors:** Alexandra Trejo-Gaytán, Axel A. Rojero-Hernández, José T. Otero-Pappatheodorou, Juan E. Gris-Gómez, Christian O. Venegas-Regin, Israel Pichardo-Casas, Andrés Gatica-Arias, José M. Villalobos-Escobedo

**Affiliations:** Tecnológico de Monterrey, Institute for Obesity Research, Monterrey, NL 64849, Mexico; University of Costa Rica, School of Biology, Plant Biotechnology Laboratory, San José, Costa Rica

**Keywords:** Fermented beverages, Genome mining, Metagenome-assembled genomes, Genome-scale metabolic modeling, Cross-feeding, Carbohydrate-active enzymes, Secondary metabolites

## Abstract

Fermentation of foods and beverages represents one of humanity’s oldest biotechnologies, generating compounds with demonstrated benefits for gut microbiota modulation, immune regulation, and metabolic health. The global rise of non-communicable chronic diseases, including obesity, type 2 diabetes, and chronic inflammation, has intensified the search for microbiome-based interventions, positioning fermented beverages as promising sources of next-generation probiotics and functional microbial consortia. Beverages such as kefir and kombucha, together with traditional Mexican fermented beverages including pozol and pulque, have been subjected to high-depth shotgun metagenomic studies generating high quality genomic resources. Systematic genomic mining efforts aimed at the functional characterization and biotechnological exploitation of these microbial communities, however, remain scarce. Here, we used a bioprospecting pipeline applied to milk-based kefir, kombucha, pozol, and pulque, integrating targeted genomic mining of genes associated with the biosynthesis of B-group vitamins, short-chain fatty acids, natural products, and CAZymes with potential to enhance starch and dietary fiber utilization upon intestinal colonization. Through genome-scale metabolic modeling of metagenome-assembled genomes, we identified microbial candidates predicted as central producers of secondary metabolites involved in pathogen control. We then used these results for the *in silico* synthetic assembly of a six-member synthetic microbial community predicted to exhibit stable cooperative growth and high metabolic functionality. Cross-feeding analysis revealed iron as one of the most widely shared elements among community members, with *Priestia flexa* from pozol, serving as a major donor of compounds involved in iron transport and as a stabilizing element within the synthetic community. This approach allows us to design a theoretical highly functional probiotic community, opening new avenues for the systematic exploitation of microbial diversity for biomedical purposes.

## Introduction

Fermented foods and beverages have been consumed across human cultures since ancestral times, representing one of the oldest and most widespread food preservation strategies (Marco *et al.,* 2021). Among the most globally popular fermented beverages today is kombucha, a drink from Eastern Asia prepared from sweetened tea inoculated with a symbiotic culture of bacteria and yeast known as SCOBY (symbiotic culture of bacteria and yeast), a starter culture (Abaci *et al.,* 2022). Kombucha is considered a functional beverage rich in bioactive compounds, antioxidants, and probiotic microorganisms, with quantifiable effects on the intestinal microbiome (Onsun *et al.,* 2025). Following fermentation, the beverage contains a diverse array of organic acids, amino acids, polyphenols, and micronutrients including B-complex vitamins and minerals such as zinc, copper, iron, and manganese, thought to underlie its health-promoting properties (Onsun *et al.,* 2025). Antimicrobial, cardioprotective, anticancer, and antidiabetic effects have also been attributed to kombucha, including reports of reduced blood glucose levels upon consumption. Although clinical evidence remains limited, the widespread global consumption of this beverage has driven scientific interest, resulting in an increasingly detailed understanding of its bioactive and healthful properties. This level of attention, however, has been largely absent for other traditionally fermented beverages with comparably rich microbial compositions but far less global visibility.

Kefir, for example, is a fermented milk-based beverage produced using kefir grains, complex microbial communities composed of lactic acid bacteria, acetic acid bacteria, and yeasts as starter culture (Tenorio-Salgado *et al.,* 2021). Current research shows that kefir consumption increases beneficial taxa such as *Lactobacillus* and *Bifidobacterium* while reducing pro-inflammatory microbes such as *Enterobacteriaceae* (Hamsho *et al.,* 2026). Kombucha and kefir beverages have expanded globally, driven by their organoleptic characteristics and perceived health benefits. The kombucha market, valued at US$4.8 billion in 2025, was projected to reach US$9.1 billion by 2033 (Grand View Research, 2026), and global markets for both kombucha and kefir have demonstrated sustained growth trajectories fueled by increasing consumer interest in digestive health and probiotic products (Chong *et al.,* 2023). This momentum is further amplified within the broader functional food sector, of which fermented beverages constitute a fast-growing segment.

Starter culture, in the context of kefir and kombucha, refers to a living microbial community, traditionally undefined at the species level, that is deliberately introduced into a substrate to drive fermentation toward a desired product, outcompeting unwanted environmental microbes through numerical dominance and rapid metabolic activity. Neither kefir grains nor the kombucha SCOBY were rationally designed. They are empirical, traditional cultures that emerged through centuries of serial back-slopping, where a portion of one successful batch is used to inoculate the next. This repeated transfer acts as an unconscious selective pressure, enriching for microbial consortia stable and well-adapted to that specific fermentation niche, without any molecular-level understanding of the community composition until the 20th century.

In Mexico, a rich pre-Columbian heritage has given rise to a diverse array of spontaneously fermented beverages that remain widely consumed today, arising not from a defined starter culture but from the microbiota naturally present in the raw substrate together with environmental microorganisms to which the substrate is exposed during fermentation. These beverages include pulque, pozol, tepache, tesgüino, sambumbia, tuba, taberna, and colonche, among others (Ojeda-Linares *et al.,* 2021). These beverages are derived from substrates such as agave sap, nixtamalized maize, cactus fruit, and palm sap, and they ferment entirely by spontaneous microbial communities shaped by local raw materials and environmental conditions. Two beverages of particular cultural and scientific relevance are pozol, a non-alcoholic maize-and cacao-based drink consumed in the southeastern Maya region of Mexico, and pulque, produced from agave aguamiel and consumed primarily in central Mexico (Supplementary Figure S1).

Using shotgun metagenomic sequencing at four key fermentation moments (0, 9, 24, and 48 h), López-Sánchez *et al*. (2023) identified 25 abundant bacterial genera throughout pozol fermentation, with *Streptococcus* as the dominant genus, accounting for 60% of all reads. Metagenome-assembled genomes (MAG) reconstruction yielded 11 high-quality bins corresponding to *Streptococcus infantarius*, *S. ferus*, *Weissella confusa*, *Lactobacillus delbrueckii*, *Limosilactobacillus fermentum*, *Lactococcus garvieae*, *Enterococcus italicus*, *Leuconostoc* sp., *Exiguobacterium* sp.*, Anoxybacillus* sp., and *Rothia* sp. Functional annotation revealed metabolic potential for starch, plant cell wall, fructan, and sucrose degradation, as well as biosynthetic modules for essential amino acids and riboflavin. The functional characterization of these MAGs, however, remained largely descriptive and was not contextualized within the broader landscape of traditional fermented beverage microbiomes. Notably, the starch-rich substrate of pozol selects for microorganisms with broad carbohydrate-active enzyme repertoires, a trait of direct relevance for probiotic applications given the capacity of such organisms to degrade complex dietary carbohydrates upon intestinal colonization.

Similarly, using whole-genome shotgun metagenomic sequencing across five stages of pulque fermentation, Chacón-Vargas *et al*. (2020) identified six key genera — *Acinetobacter, Lactobacillus*, *Lactococcus, Leuconostoc, Saccharomyces,* and *Zymomonas* — whose relative abundances shifted dynamically depending on sucrose consumption and concurrent increases in ethanol and lactic acid production. Predictive functional profiling revealed enrichment of folate biosynthesis genes in mature pulque across all major bacterial genera, consistent with the recognized nutritional value of this beverage. No MAGs were reconstructed, however, leaving the metabolic potential of individual community members unresolved at the genomic level.

In this work, we reanalyzed publicly available high-depth shotgun metagenomic data from kombucha, kefir, pozol, and pulque, reconstructing 77 MAGs and performing metabolic pathway enrichment and enzymatic function analyses to enable a genome-resolved comparison across these four traditionally fermented beverages (Fig. S1). Kombucha-associated genomes show marked enrichment in complete vitamin B12 biosynthesis pathways alongside short-chain fatty acids (SCFA) production capacity, the pozol microbiome is enriched in CAZymes with biosynthetic potential for vitamins B9 and K, and *Priestia flexa* from pozol emerges as a high-value probiotic candidate combining broad carbohydrate degradation capacity with SCFA production and as an important iron donor within the microbial community. In pulque, *Hafnia alvei* stands as a promising B12-producing strain.

To move beyond descriptive genomics, we reconstructed genome-scale metabolic models (GEMs) from the 13 highest-quality MAGs and performed community flux balance analysis using microbial community modeling (MICOM v0.39.0; Diener *et al.,* 2020) to evaluate the metabolic connectivity and cooperative stability of candidate synthetic communities. Through compatibility scoring, metabolic exchange network analysis, and leave-one-out inducer-repressor profiling, we identified a six-member synthetic microbial community (SynCom) predicted to sustain stable cooperative growth while preserving key probiotic traits, including SCFA production, vitamin biosynthesis, and complex carbohydrate degradation.

The objective of this study was to determine whether genome-resolved metagenomics and metabolic modeling could be integrated to identify and rationally design candidate probiotic synthetic communities from traditional fermented beverages.

## Methods

Shotgun metagenomic sequencing data from four fermented beverages, kombucha (PRJNA833075), kefir (PRJNA704713), pozol (PRJNA64886), and pulque (PRJNA603591) were retrieved from publicly available datasets in the NCBI Sequence Read Archive. All sequencing runs were downloaded in FASTQ format using the SRA Toolkit. Metadata for each run, including accession numbers, sequencing platform, library strategy, and associated publication, are provided in Supplementary Table S1. In addition, Supplementary Table S2 further provides geographic detail.

### Read quality control and trimming

Raw sequencing reads were assessed for quality using FastQC v0.12.1 (Andrews, 2010). Adapter sequences and low-quality bases were removed using Trimmomatic v0.40 (Bolger *et al.,* 2014) in paired-end mode with the following parameters: ILLUMINACLIP TruSeq3-PE.fa, LEADING 3, TRAILING 3, SLIDINGWINDOW 4:15, and MINLEN 36. Only read pairs in which both mates passed quality filtering were retained for downstream analysis.

### Taxonomic classification and diversity analysis

Taxonomic classification of trimmed reads was performed using Kaiju v1.10.1 (Menzel *et al.,* 2016) with the NCBI BLAST non-redundant protein database in nr+euk mode, which enables protein-level classification and improves sensitivity for divergent or partially sequenced taxa. Output files were converted to taxonomic profiles at the genus level using kaiju2table and further processed for visualization (Supplementary Table S3). Alpha diversity metrics (observed richness, Shannon index, and Simpson index) and beta diversity (Bray–Curtis dissimilarity) were computed in R using the tidyverse v2.0.0 (Wickham *et al.,* 2019) and vegan v2.7-5 (Oksanen *et al.,* 2022) packages. Non-metric multidimensional scaling ordination was used to visualize community structure. Significance of inter-group differences was tested using the vegdist function from vegan v2.7-5. Genus-level relative abundance was visualized with ggplot2 v4.0.3 (Wickham, 2016).

### Metagenomic assembly and binning

Trimmed reads from each metagenome were assembled *de novo* using metaSPAdes v4.1.0 (Nurk *et al.,* 2017) with default k-mer sizes and the `--meta` flag. Prior to binning, trimmed reads were mapped back to their respective assemblies using Bowtie2 v2.5.5 (Langmead & Salzberg, 2012) to generate per-contig coverage profiles. Binning was performed with MetaBAT2 v2.18 (Kang *et al.,* 2019) and MaxBin2 v2.2.7 (Wu *et al.,* 2016) independently, and the resulting bins were refined and dereplicated using DAS_Tool v1.1.6 (Sieber *et al.,* 2018), which selects the highest-quality non-redundant set of bins from multiple binners. Contigs shorter than 1,000 bp were excluded prior to binning. The resulting MAGs were dereplicated across all beverages at 95% average nucleotide identity (ANI) using dRep v3.6.2 (Olm *et al.,* 2017) to avoid redundancy in downstream analyses.

### MAG quality assessment

The completeness and contamination of all MAGs were evaluated using Benchmarking Universal Single-Copy Orthologs (BUSCO v6.1.0) (Manni *et al.,* 2021) with the bacteria_odb10 lineage dataset, which provides a curated set of single-copy orthologs conserved across bacteria (Supplementary Table 4). Only MAGs with completeness ≥ 75% and contamination ≤ 10% were retained for downstream analyses, following accepted quality thresholds for medium-to high-quality MAGs. MAGs with completeness ≥ 95% were classified as high quality. We used BUSCO for completeness assessment because it is the only tool capable of scoring both prokaryotic and eukaryotic genomes within a single framework of near-universal single-copy orthologs (Tegenfeldt *et al*., 2025). This was essential here, as our dataset includes eukaryotic genomes, the yeast *Brettanomyces bruxellensis* and the maize chloroplast genome from pozol (eukaryota_odb10), with tools restricted to prokaryotic markers such as CheckM2 v1.1.0 cannot assess.

### Taxonomic MAG classification

MAGs were taxonomically classified using the Genome Taxonomy Database Toolkit (GTDB-Tk v2.7.1/2.7.2, Chaumeil *et al.,* 2022) with the GTDB r214 reference database, which applies a standardized phylogenetic framework based on concatenated marker gene trees and ANI comparisons. Each MAG was assigned to the lowest possible taxonomic rank; those that could not be classified to the species level were retained in the dataset as unclassified MAGs, as they may represent undescribed lineages of ecological relevance (Supplementary Table 5).

### Targeted functional enrichment of health-relevant pathways

To analyze the functional potential of beverage microbiomes for health-relevant processes, annotated gene sets were classified in three major functional categories: complex carbohydrate degradation (CAZymes), short-chain fatty acid (SCFA) biosynthesis and metabolism (Supplementary Table 6), and vitamin biosynthesis (B9, B12, and K) (Supplementary Table 7). Supplementary Table S6 provides target gene lists for each subcategory were assembled from KEGG pathway maps (Kanehisa *et al.,* 2023), gene-name patterns, annotation keyword searches, CAZy/dbCAN family annotations and curated literature. CAZyme-associated genes were identified using both eggNOG-mapper v2.1.x annotations (Cantalapiedra *et al*., 2021) and dbCAN4 (Zhang *et al.,* 2018; Zheng *et al.,* 2023) with HMMER against the CAZy HMM profiles (Drula *et al.,* 2022). For each sample, target gene counts per subcategory were calculated and normalized to the total number of annotated genes to obtain relative abundances. Subcategory-level differences across beverages were tested with the Kruskal–Wallis test, and pairwise comparisons were performed with Dunn’s test with Benjamini–Hochberg correction (p-adjusted < 0.05). Only subcategories with a significant global test and at least one significant pairwise comparison were retained for visualization. CAZyme guild classification was performed by assigning each MAG to a guild based on its dominant substrate-specific enzyme profile and repertoire breadth. Scatterplot analyses of CAZyme versus SCFA targets, and of vitamin biosynthesis versus consumption targets, were used to identify functionally integrative MAGs across beverages.

### Biosynthetic gene cluster prediction

Biosynthetic gene clusters (BGCs) in each MAG were predicted using antiSMASH v8.0.2 (Blin *et al.,* 2023) with the `--genefinding`-tool prodigal and `--fullhmmer` flags enabled for maximal sensitivity. All BGC antiSMASH-supported classes were considered, including polyketide synthases, non-ribosomal peptide synthetases, terpenes, bacteriocins, siderophores, and others. Each predicted BGC was characterized by class, the predicted core compound identity, and the similarity score to the closest known BGC in the MIBiG reference database (Terlouw *et al.,* 2023), which was used to distinguish high-confidence, medium-confidence, and low-confidence predictions.

### MAG-level CAZyme guild classification and substrate specialization

To resolve carbohydrate-degradation potential at the genome level, the six CAZyme subcategories (starch, cellulose/cellobiose, hemicellulose/xylan, fructan/inulin, pectin/oligosaccharides, and other glycoside hydrolase families) were tabulated per MAG, and each MAG was assigned to a functional guild based on its dominant subcategory and repertoire breadth. MAGs that had at least five of the six subcategories with no single subcategory exceeding 50% of total CAZyme targets were classified as broad-spectrum degraders, and the remaining MAGs were assigned to the guild corresponding to their dominant subcategory (starch-enriched, fructan/inulin-enriched, plant cell wall-associated, cellulose/cellobiose-rich, or other-CAZyme-dominant). Substrate specialization was quantified per MAG as the fraction of CAZyme targets assigned to starch and to fructan/inulin, and the distribution of these scores across beverages was compared with the Kruskal–Wallis rank-sum test; the starch specialization score differed significantly among beverages (p = 1.4 × 10⁻⁷), being highest in pozol. The pozol MAGs with the highest starch specialization scores, restricted to genomes with at least eight total CAZyme targets, were displayed as a target-level count heatmap to identify the enzyme families driving the starch signal. To assess co-occurrence of carbohydrate-degradation and fermentative potential within genomes, total CAZyme-associated targets and total SCFA-related targets were tabulated per MAG and compared in a scatterplot, with MAGs exceeding the median of both axes identified as candidate functional bridge organisms.

### Statistical analysis of biosynthetic pathway and CAZyme distributions

All statistical comparisons of target-gene counts and relative abundances across beverages were performed in R. Given the non-normal distribution of genomic functional data, non-parametric tests were used throughout. Global differences among the four beverage groups were assessed using the Kruskal–Wallis rank-sum test. Significant global results (p < 0.05) were followed by pairwise Dunn’s tests, with p-values adjusted for multiple comparisons using the Benjamini–Hochberg false discovery rate (FDR) method. Results were considered statistically significant at FDR-adjusted p < 0.05. For the BGC and CAZyme enrichment analyses, counts per MAG were compared across beverages using the same two-step framework. Visualizations were produced using ggplot2 (Wickham, 2016), ggpubr v0.6.3 (Kassambara, 2023), and ComplexHeatmap v2.26.1 (Gu, 2022) in R.

### Genome-scale metabolic model reconstruction

GEMs were reconstructed from MAG nucleotide assemblies using CarveMe v1.6.6 (Machado *et al.,* 2018) with the `--dna` flag, which performs gene prediction, functional annotation via DIAMOND v2.1.8 (Buchfink *et al.,* 2021) against the BiGG/AGORA database (King *et al.,* 2016; Magnúsdóttir *et al.,* 2017), and model carving in a single workflow. Each GEM was validated by performing flux balance analysis (FBA) using COBRApy v0.26 (Ebrahim *et al.,* 2013), confirming that all models achieved optimal solutions. Model quality metrics including number of reactions, metabolites, genes, and exchange reactions were recorded for each GEM (Table S1).

### Community metabolic modeling

Community flux balance analysis was performed using MICOM v0.39.0 (Diener *et al.,* 2020). All 13 GEMs were integrated into a community model with equal relative abundances (1/n per member). Community growth was optimized using the cooperative tradeoff algorithm at a tradeoff fraction of 0.5, which balances individual and community-level growth maximization. Exchange fluxes were extracted for all community members and used for downstream interaction analysis.

### Metabolic exchange network analysis

Exchange reaction fluxes were filtered to remove stoichiometric artifacts (H₂O, H⁺, CO₂, phosphate, ammonium, O₂), and interactions with significance below 0.05% of total organismal flux were excluded. Remaining interactions were classified into five metabolite categories: amino acids, carbon sources, cofactors, nucleotides, and ions, based on BiGG metabolite identifiers. Directed exchange networks were constructed using NetworkX v3.x (Hagberg *et al.,* 2007) and visualized using Matplotlib v3.x (Hunter, 2007), with edge weights proportional to exchange flux magnitude and node colors reflecting compatibility scores.

### Compatibility scoring

A composite compatibility score was computed for each genome based on three normalized metrics. (1) net metabolic contribution (exported flux minus imported flux, weight 0.50), (2) inter-organismal connectivity (sum of unique producer and consumer partners, weight 0.30), and (3) metabolite diversity (total unique metabolites exchanged, weight 0.20). Weights were assigned in decreasing order of biological priority. Net contribution was weighted highest as the primary determinant of donor-receiver balance, connectivity second as a measure of network integration, and diversity lowest as a secondary indicator of functional breadth. Each metric was min-max normalized to the interval [0,1] across the community before weighted summation. Genomes with compatibility scores below 0.35 were considered low-compatibility candidates for exclusion from the SynCom.

### Inducer-repressor analysis

To identify induction and repression relationships, a leave-one-out (LOO) analysis was performed following the approach by Zelezniak *et al*. (2015). For each genome, a reduced community model was constructed excluding that genome, and cooperative tradeoff FBA was repeated under identical conditions using MICOM. The percent change in growth rate of each remaining member relative to the full community baseline was calculated as:

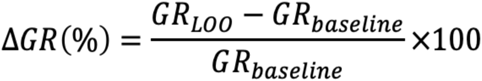

Genomes were classified as inducers of a target if their removal caused a growth rate decrease exceeding 5% (Δ < −5%), and as repressors if their removal caused a growth rate increase exceeding 5% (Δ > +5%). Interactions within ±5% were classified as neutral.

### SynCom design

The final SynCom was assembled by integrating three criteria: (1) compatibility score above the 0.35 threshold, (2) inducer status, genomes whose removal significantly reduced community growth were prioritized for inclusion, and (3) donor-receiver balance, preference was given to net metabolic donors. The resulting six-member SynCom was validated by re-running the full community simulation pipeline independently to confirm stable cooperative growth and retention of key metabolic interactions (Supplementary Table 8 & 9).

## Results

### Taxonomic structure and diversity distinguish spontaneously fermented from starter culture-based beverages

We first analyzed the taxonomic composition and diversity of the fermented beverages using samples from different geographical regions (Fig. 1A; Fig. S1). Genus-level profiling revealed marked differences in microbial community structure across beverages, with each fermentation system harboring distinct microbial assemblages. Despite this separation, several genera were consistently detected across multiple beverages, suggesting the existence of shared microbial components with conserved functional roles.

**Figure 1.**
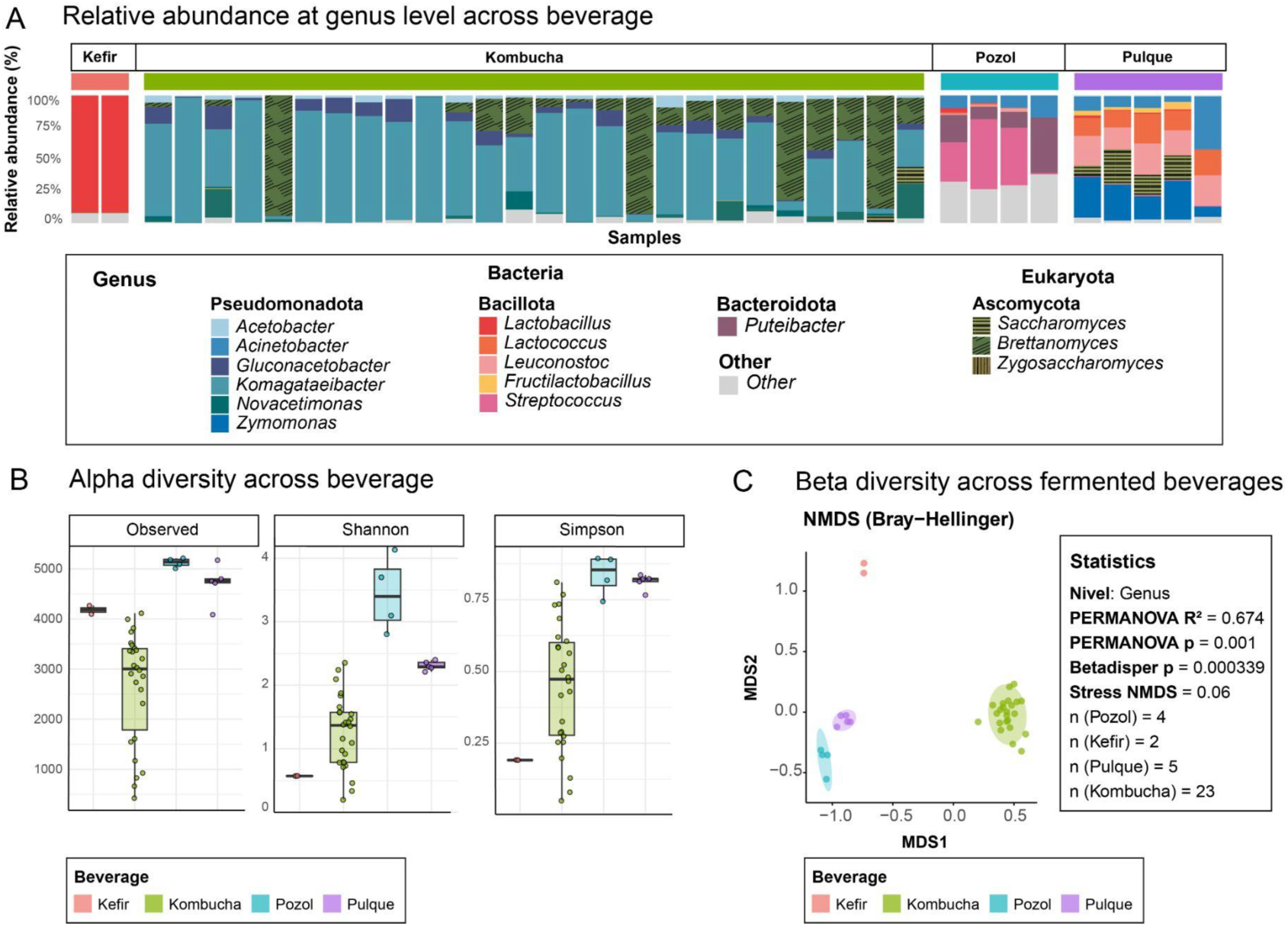
Microbiological analysis and diversity of fermented beverages. (**A**) Microbial composition at the genus level in kefir, kombucha, pozol, and pulque samples, highlighting both intra-and inter-beverage variability. (**B**) Alpha diversity (richness, Shannon, and Simpson indices) across fermented beverage types. Each point represents an individual sample. (**C**) Beta diversity analysis, using non-metric multidimensional scaling ordinations constructed from Bray–Curtis dissimilarities on Hellinger-transformed relative abundances (“Bray-Hellinger”), shows clear structural separation among microbial communities associated with each beverage type. Community composition differed significantly among beverages (PERMANOVA, *R*² = 0.674, *p* = 0.001), although multivariate dispersion also differed among groups (beta dispersion, *p* = 0.000339). Stress = 0.06. Sample sizes were for kefir (n = 2), kombucha (n = 23), pozol (n = 5), and pulque (n = 6).

Traditional beverages, particularly pozol and pulque, displayed consistently higher microbial alpha diversity than kefir and kombucha, most notably in the Shannon and Simpson indexes, which capture both richness and evenness (Fig. 1B). This pattern is consistent with their spontaneous fermentation from complex substrates such as maize and agave sap, where raw materials and the local environment shape open, diverse microbial communities rather than a single defined starter inoculum. Kombucha showed comparable Observed richness to pozol and pulque in some samples, but consistently lower Shannon and Simpson values, and a wide range of variation across samples, suggesting that its consortium is dominated by a small number of taxa (mainly acetic acid bacteria and yeasts) rather than lacking taxa altogether. Kefir showed the lowest diversity across all three indexes despite comparable richness to pulque, reflecting strong dominance by its defined grain-associated consortium. Together, these results indicate that the open, substrate-driven fermentations of pozol and pulque support richer and more even microbial communities than the starter-driven fermentations of kefir and kombucha.

Beta diversity analysis reinforced these observations, showing clear sample clustering by beverage type (Fig. 1C), indicating strong ecological structuring across fermentation systems. Despite sharing the broad category of fermented beverages, genus-level composition revealed minimal taxonomic overlap among beverages, suggesting that each fermentation system promotes a highly substrate-specific microbial assemblage. The consistency of community composition within samples of the same beverage, combined with its distinctiveness across beverage types, points to stable and long-standing co-fermentation histories shaped by the physicochemical properties of each substrate, the selective pressures imposed by fermentation conditions, and potentially by centuries of human-guided domestication of these microbial communities.

### High-quality genome assembly from the metagenomes of the four fermented beverages

Seventy-seven MAGs were recovered across the four beverages: kombucha (24), pozol (25), pulque (24), and kefir (4). All MAGs displayed completeness above 75% with low contamination and fragmentation levels as assessed by BUSCO (Fig. 2A). A subset reached completeness above 95%, reflecting high genomic integrity and supporting their use in high-resolution comparative and functional analyses.

**Figure 2.**
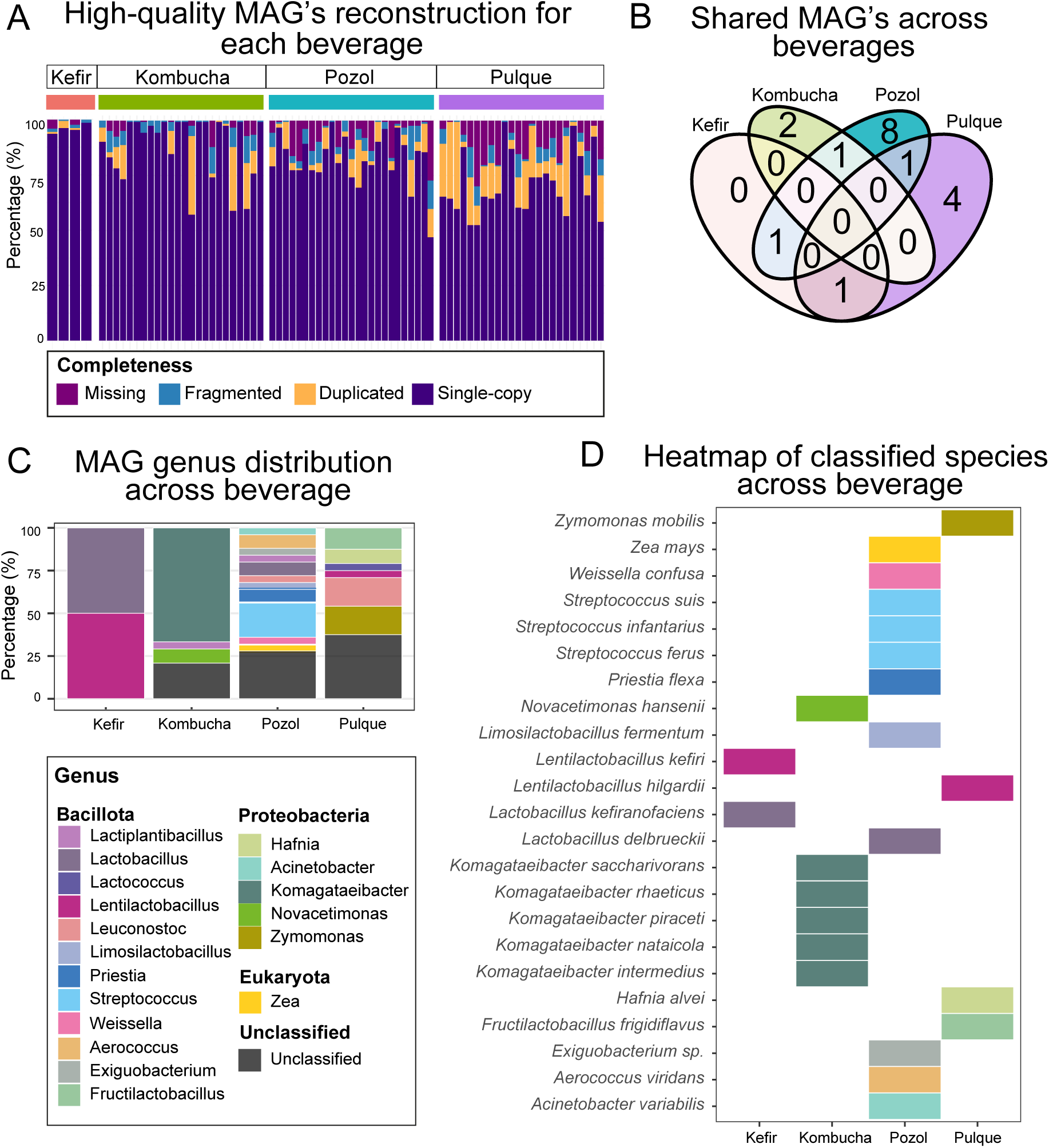
Genome quality, taxonomic distribution, and shared microbial features across fermented beverages. (**A**) Completeness and redundancy of reconstructed metagenome-assembled genomes (MAGs) assessed using BUSCO, with completeness greater than 75%, indicating high genome completeness with low levels of duplication and fragmentation. (**B**) Shared high quality reconstructed genomes between beverages. (**C**) Relative abundance of the number of MAGs in each genus at taxonomic distribution. (**D**) Heatmap showing the distribution of MAGs across individual samples (completeness > 95%), where each cell represents the number of MAGs assigned to a given species in each beverage, revealing distinct and beverage-specific microbial compositions.

Taxonomic distribution of the recovered MAGs revealed clear microbial community structuring across beverages (Fig. 2B). Kombucha was dominated by MAGs belonging to the genus *Komagataeibacter,* whereas pulque and pozol showed broader taxonomic diversity with well-represented genera spanning multiple lineages, consistent with the more complex microbial communities characteristic of spontaneous fermentation. Notably, unclassified MAGs with high genomic integrity were recovered from pozol, kombucha, and pulque, suggesting the presence of previously unreported species that may play relevant roles in these fermentation systems.

Genus-level overlap analysis revealed that *Lactobacillus* and *Lentilactobacillus* were the only genera shared across beverages, though with distinct distributions (Fig. 2C). *Lactobacillus* was present in both kefir and pozol, while *Lentilactobacillus* was shared between kefir and pulque. All remaining genera showed restricted distributions, reflecting specific ecological adaptations to each fermentation environment. Of particular interest, *Zymomonas*, a genus widely associated with fermentation processes (Weir *et al.,* 2016), was detected exclusively in pulque. The recovery of high-integrity unclassified MAGs across three of the four beverages points to unexplored metabolic diversity and the potential for discovering novel functional pathways and bioactive compounds within these traditional fermentation systems.

### Distribution and annotation confidence of biosynthetic gene clusters across fermented beverages

To assess the secondary metabolite production potential across beverages, we predicted BGCs from MAGs recovered from kombucha, pozol, and pulque (Fig. 3A and 3B). No BGCs were predicted in any of the kefir-associated MAGs, consistent with the dominance of lactic acid bacteria in this fermentation system. Kombucha harbored the highest number of predicted BGCs, with *Komagataeibacter rhaeticus* encoding up to 5 BGCs, most at high similarity, followed by other *Komagataeibacter* species and *Brettanomyces bruxellensis* with 2 to 3 BGCs each (Fig. 3A). In pozol, an unclassified organism carried two high-similarity BGCs, while *Priestia flexa* and *Lactococcus garvieae* contributed additional clusters at medium or low confidence. Pulque showed the lowest overall confidence, with *Lactococcus piscium* encoding the most BGCs (2), though largely at low similarity.

**Figure 3.**
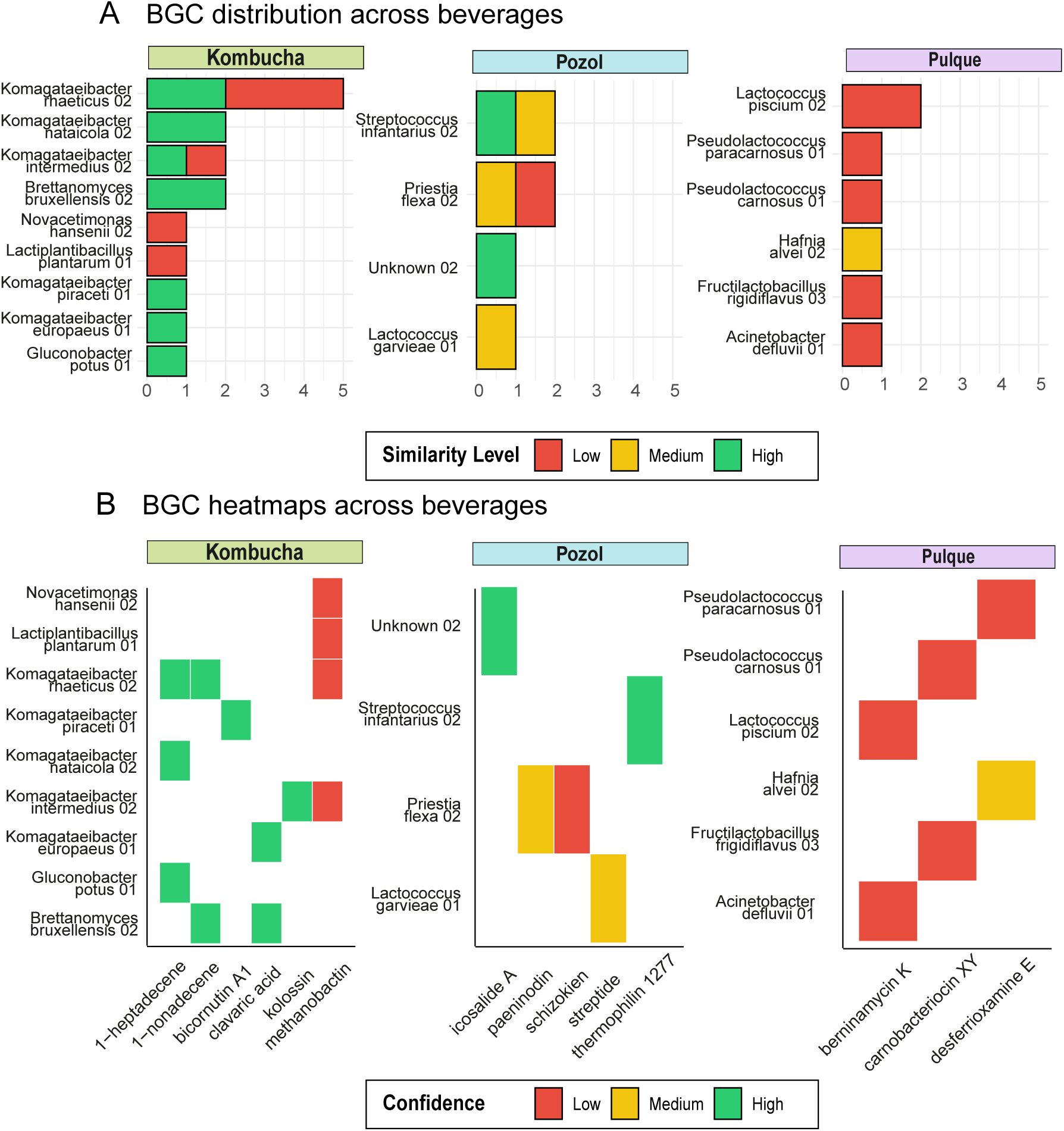
Biosynthetic gene cluster (BGC) content analysis of MAGs from fermented beverages. (**A**) BGC distribution counts in kombucha, pozol, and pulque. The x axis represents the total number of BGCs detected per species. (**B**) Heatmaps displaying the predicted compounds encoded by BGCs identified across the three beverages. The x axis shows the different detected compounds as predicted by antiSMASH, while the y axis shows the assembled species for each beverage. Colors indicate antiSMASH similarity levels: red = low, yellow = medium, and green = high.

The compound-level heatmap (Fig. 3B) confirmed that kombucha BGCs encoded the most reliably predicted products, including 1-heptadecene and 1-nonadecene. In pozol, the high-confidence predictions were icosalide A from the unclassified organism and thermophilin 1277 from *Streptococcus infantarius*. In pulque, desferrioxamine E is predicted with medium confidence in *H. alvei*. *Priestia flexa*, in turn, harbored a BGC with predicted similarity to icosalide A, an unusual two-tailed lipocyclopeptide depsipeptide with demonstrated antibacterial activity including inhibitory action against *Mycobacterium tuberculosis* at MIC values between 2 and 10 μg mL⁻¹ (Dangi *et al*., 2023). The presence of BGCs encoding compounds with distinct antibacterial mechanisms across two co-occurring pozol organisms suggests that this traditionally consumed beverage harbors an underexplored reservoir of antimicrobial biosynthetic diversity.

### The genomic content of each beverage encodes differentially abundant functions in health-relevant processes

To identify functional differences across the four fermented beverage microbiomes, we compared annotated target-gene subcategories and retained only those showing a significant Kruskal-Wallis test and at least one significant Dunn pairwise comparison (Fig. 4). This approach highlighted three major functional axes distinguishing the beverages: complex carbohydrate degradation, SCFA metabolism, and vitamin biosynthesis.

**Figure 4.**
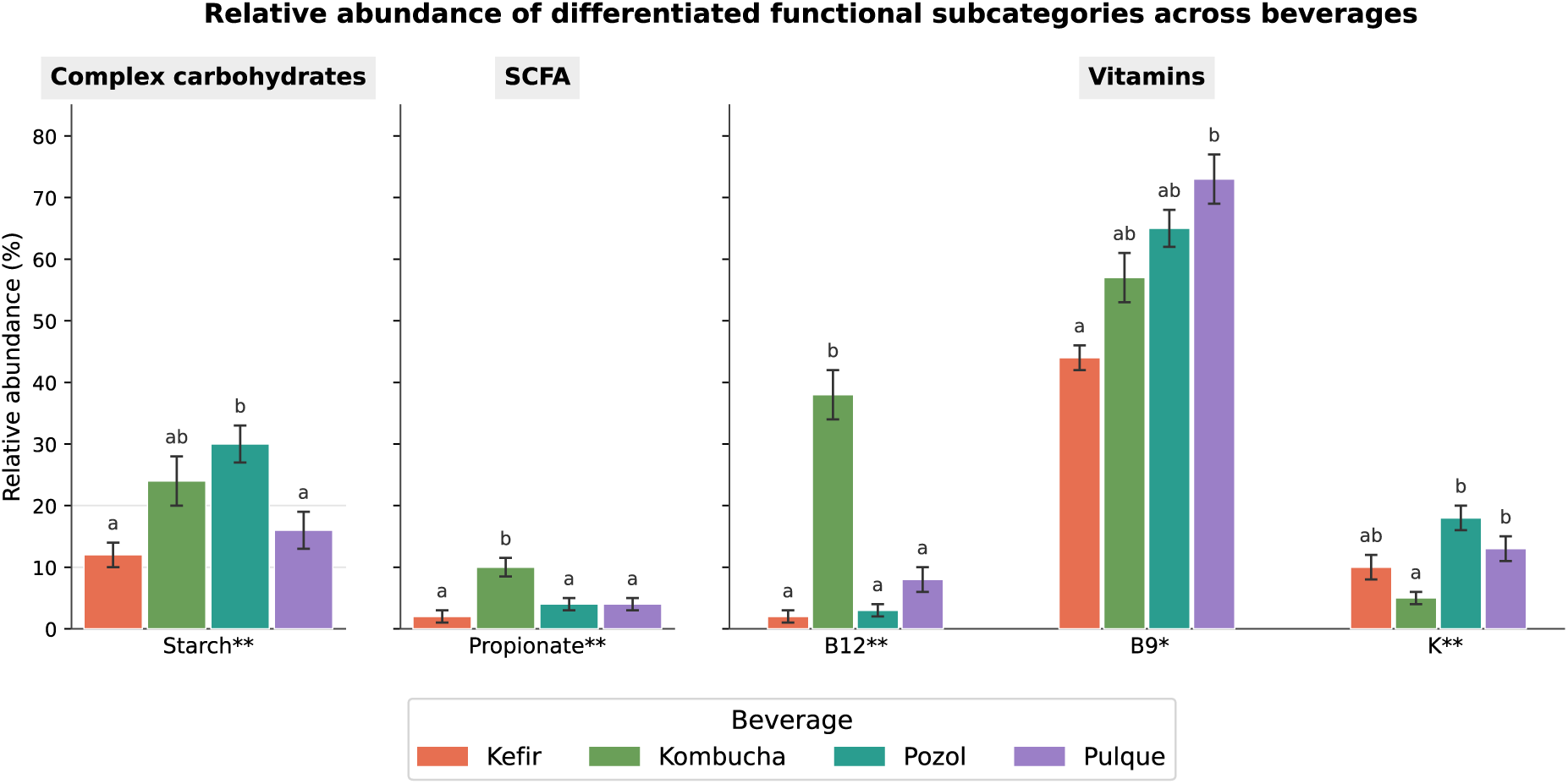
Global screening of robustly differentiated functional subcategories across fermented beverages. Vertical bar plots display the relative abundance of annotated target genes (%) for functional subcategories that passed a two-step statistical filter: a significant global Kruskal-Wallis test followed by at least one significant Dunn pairwise comparison. Results are organized into three major functional categories: complex carbohydrates, short-chain fatty acids (SCFA), and vitamins. Each bar represents the mean relative abundance for a given beverage, kefir (red), kombucha (green), pozol (blue), and pulque (purple), with error bars. Letters above bars denote statistically significant pairwise differences based on Dunn’s post-hoc test, where groups sharing the same letter are not significantly different. Asterisks next to subcategory names indicate the level of significance of the global Kruskal-Wallis test (* p < 0.05, ** p < 0.01).

Within complex carbohydrate degradation, starch-related genes were most abundant in pozol, significantly higher than in kefir, consistent with its maize-based substrate. Kombucha and pulque occupied intermediate positions with no significant difference between them. Within SCFA metabolism, propionate-related genes were most abundant in kombucha.

The most pronounced differences were observed in vitamin biosynthesis. Vitamin B12 biosynthesis genes were markedly enriched in kombucha relative to other beverages, consistent with the high B-vitamin content previously reported for this beverage. Folate (B9) biosynthesis was broadly distributed but higher in pulque than in the other three beverages. Vitamin K biosynthesis was significantly depleted in kombucha relative to the other beverages. These findings indicate that each beverage harbors a distinct biosynthetic repertoire for B-group vitamins, micronutrients directly relevant to hematopoiesis, neural function, and coagulation. Taxon-level analyses further revealed the distribution of these vitamin-related pathways among dominant microbial taxa (Supplementary Figure S2).

Folate biosynthesis represented the predominant vitamin synthesis pathway among most taxa from kefir, pozol, and pulque, whereas kombucha-associated microorganisms showed a higher relative contribution of cobalamin biosynthesis within their vitamin metabolism repertoire (Supplementary Figure S2A). In contrast, vitamin uptake profiles were dominated by B12 transport pathways across taxa from all beverages, suggesting that cobalamin acquisition is a widespread strategy among fermented beverage microorganisms, including taxa lacking complete biosynthetic pathways (Supplementary Figure S2B). Consistent with this pattern, B12 biosynthesis potential was mainly restricted to a limited number of kombucha-associated taxa, including *Komagataeibacter* spp., *Gluconobacter potus*, and *Brettanomyces bruxellensis*, whereas B12 uptake genes were broadly distributed among microbial taxa from all fermented beverages (Supplementary Figure S3).

Although the most prominent differences involved carbohydrate metabolism, SCFA production, and vitamin biosynthesis, additional functional enrichments were also identified in Clusters of Orthologous Groups (COG) categories related to stress response, ion transport, secondary metabolism, intracellular trafficking, and defense mechanisms (Supplementary Figure S4), further supporting the distinct ecological and metabolic specialization of each beverage microbiome.

### MAG-level CAZyme and SCFA profiling identifies functional bridge organisms across beverages

Based on the results above, we sought to identify which specific MAGs exhibit properties found at the global level. To characterize the carbohydrate-active potential encoded by the assembled MAGs, we grouped CAZyme-associated targets into substrate-specific categories expressed as a percentage of total CAZyme targets per beverage (Fig. 5A). Cellulose/cellobiose-associated targets were the most broadly shared component across all beverages, suggesting a conserved carbohydrate-active background. Beyond this shared baseline, beverage-specific patterns were clear. Pozol showed the highest proportion of starch-associated targets, consistent with its maize-based substrate; pulque was enriched in fructan/inulin targets reflecting agave-derived carbohydrates; and kombucha displayed a broad CAZyme profile consistent with SCOBY-associated polysaccharide turnover.

**Figure 5.**
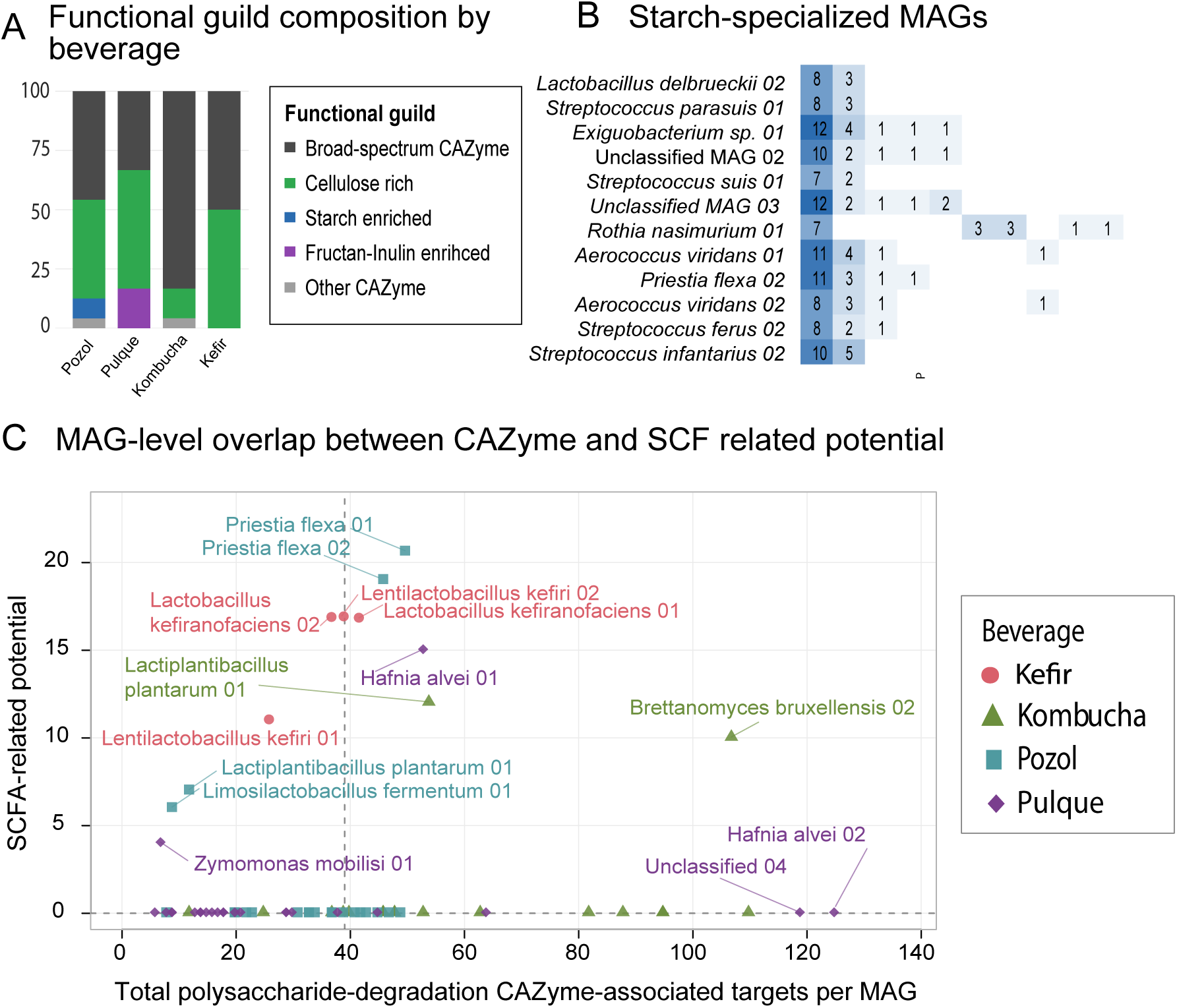
Polysaccharide-degradation potential and SCFA-linked metabolic capacity across fermented beverage MAGs. (**A**) Stacked bar plot showing the relative composition of CAZyme-associated functional profiles per beverage. Each bar represents the percentage of MAGs assigned to each CAZyme functional category, including broad-spectrum CAZyme, cellulose/cellobiose-rich, starch-enriched, fructan/inulin-enriched, plant cell wall-associated, and other CAZyme-dominant intermediate profiles. (**B**) Heatmap displaying the substrate-specific CAZyme target counts for the top-ranked MAGs per beverage, selected by their starch specialization score in pozol and fructan/inulin specialization score in pulque. Each cell represents the number of targets assigned to a given enzyme family or target gene for a given MAG. (**C**) Scatterplot comparing the total number of CAZyme-associated targets against the total number of SCFA-related targets for each MAG. Each point represents a single MAG colored by beverage of origin. Selected MAGs with high SCFA-related potential or combined CAZyme–SCFA signals are labeled, highlighting potential functional bridge organisms that may link matrix polysaccharide degradation to downstream fermentative metabolism.

At the MAG level, guild classification showed that degradation potential is unevenly distributed across communities (Fig. 5A). The starch signal in pozol was driven by enzyme families GH13, K01176, pulA, amyA, GH57, GH15, malP, glgX, and K01214, distributed across MAGs assigned to *Exiguobacterium* sp. AT1b, *Aerococcus viridans*, *Streptococcus* spp., *Rothia nasimurium*, *Lactobacillus delbrueckii*, *Priestia flexa*, and unclassified organisms (Fig. 5B).

When we examined whether MAGs with high CAZyme potential also encoded SCFA-related capacity, the relationship proved non-linear; most MAGs carried either CAZyme or SCFA targets but not both, indicating that these functions are largely distributed across different community members (Fig. 5C). A subset of MAGs did bridge both capacities, and among these *Priestia flexa* was the most functionally integrated, combining the highest CAZyme target counts in pozol with elevated SCFA-related potential within a single genome. Other MAGs showing combined CAZyme and SCFA signals included *Lactobacillus kefiranofaciens, Lentilactobacillus kefiri, Hafnia alvei, Lactiplantibacillus plantarum, Brettanomyces bruxellensis, Limosilactobacillus fermentum,* and *Zymomonas mobilis,* supporting the notion of metabolic specialization and cross-functional connectivity within these fermented beverage microbiomes. Interestingly, a pair of unclassified MAGs from pulque were also predicted as highly efficient polysaccharide degraders, potentially serving as donors of simple carbohydrate sources for other strains.

Notably, *P. flexa* 01 and 02, a MAG recovered from pozol, exhibits a striking combination of high SCFA biosynthetic gene content and broad CAZyme repertoire, positioning it as a compelling probiotic candidate. Remarkably, its dual metabolic potential appears to rival or even surpass that of well-established kefir probiotics such as *Lactobacillus kefiranofaciens* and *Lentilactobacillus kefiri*, suggesting that pozol harbors underexplored microbial taxa with exceptional functional capacity. A second organism, *H. alvei* 01, recovered from pulque, also displays a noteworthy profile, combining a diverse CAZyme arsenal with SCFA biosynthetic potential, indicating a possible role in complex carbohydrate degradation coupled to SCFA production within the fermentation community.

### Vitamin biosynthesis potential across fermented beverage MAGs reveals strain-level net producers as key functional contributors

To resolve which MAGs contributed most to vitamin biosynthesis across beverages, we compared the proportion of biosynthesis versus consumption-related targets per beverage (Fig. 6A). Kombucha and pozol showed a higher biosynthesis-to-consumption ratio than kefir and pulque, which displayed more balanced or consumption-leaning profiles.

**Figure 6.**
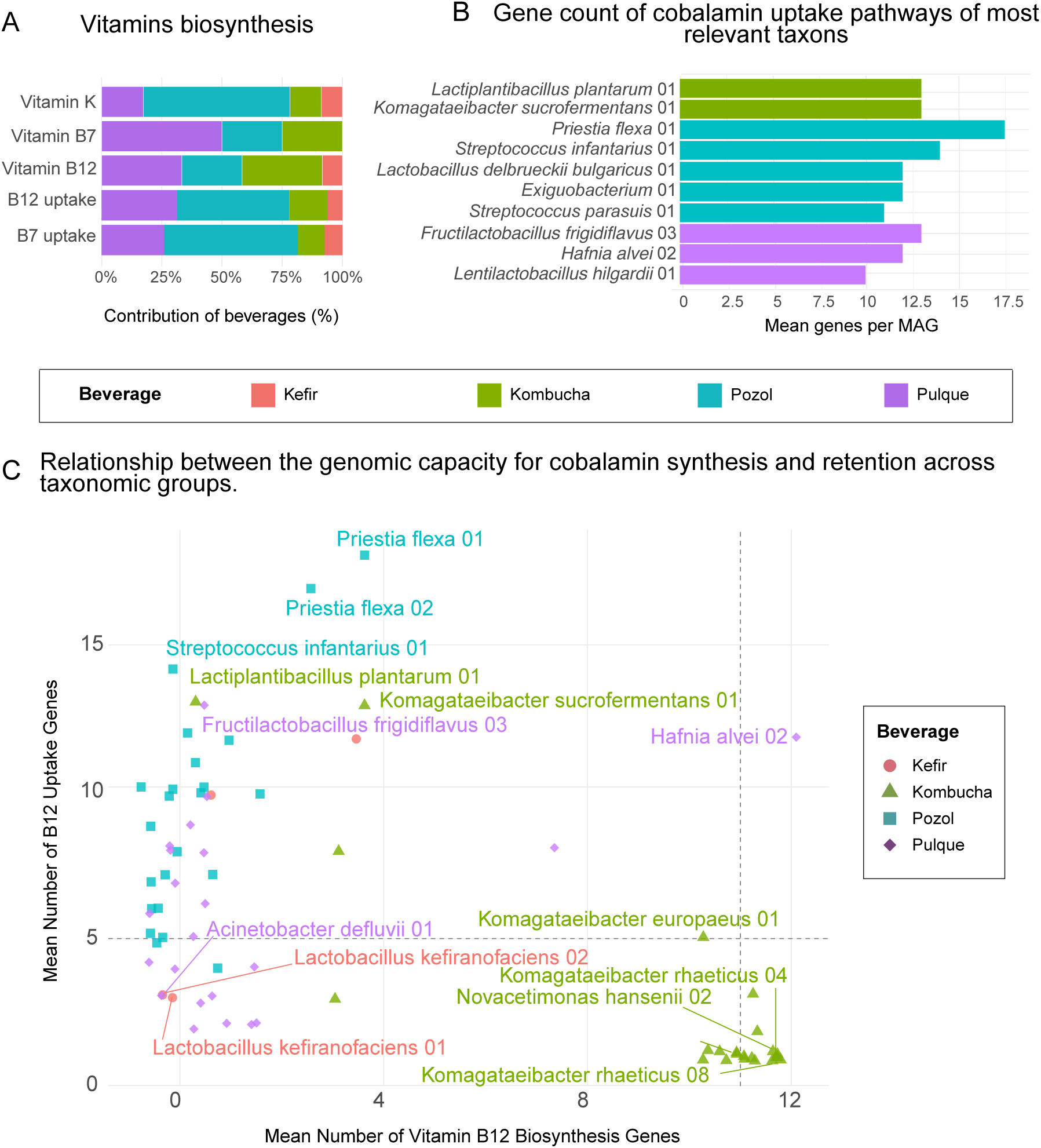
Vitamin B12 biosynthesis and uptake potential of dominant taxa across fermented beverages. (**A**) Stacked horizontal bar charts show the percentage contribution of each fermented beverage to the predicted biosynthesis and uptake potential of vitamin B12-related pathways, from 0– 100%. (**B**) Paired horizontal bar charts display the mean number of genes per MAG associated with cobalamin biosynthesis and B12 uptake across the most relevant taxa. The biosynthesis panel highlights *Komagataeibacter* spp. as the dominant contributors, with strains encoding around 10 or more biosynthesis-related genes, while the uptake panel reveals a broader taxonomic distribution of uptake capacity across beverages, with each bar colored by beverage of origin. (**C**) Four-quadrant scatterplot where each point represents a distinct MAG plotted according to its mean number of vitamin B12 biosynthesis genes on the x axis and its mean number of B12 uptake genes on the y axis. Points are colored by beverage and dashed lines delineate quadrants. Selected MAGs of functional interest are labeled.

The top biosynthesis contributors differed markedly between beverages. In kombucha, multiple *Komagataeibacter* species including *K. piraceti, K. nataicola, K. europaeus, K. intermedius,* and *K. rhaeticus* ranked highest in vitamin biosynthesis targets (Fig. 6B). In pozol and pulque, the leading contributors were *Lactiplantibacillus plantarum*, *Priestia flexa*, *Streptococcus infantarius, Lactobacillus delbrueckii bulgaricus, Fructilactobacillus frigidiflavus,* and *Hafnia alvei* (Fig. 6C).

Plotting biosynthesis against consumption targets for each MAG (Fig. 6D) allowed us to identify net producers from net consumers. *Priestia flexa* stood out with high biosynthesis and low consumption, reinforcing its role as a multifunctional keystone organism in pozol, a finding that complements its identification as a CAZyme and SCFA bridge in the previous analysis. *Streptococcus infantarius, Fructilactobacillus frigidiflavus,* and *Lactiplantibacillus plantarum* also showed net producer profiles. By contrast, *Hafnia alvei* and *Komagataeibacter nataicola* showed both high biosynthesis and high consumption, suggesting they internally recycle much of what they produce. Strains encoding vitamin biosynthesis without proportionally high consumption represent the most promising candidates for functional enrichment of fermented foods, as they are likely to contribute vitamins to the final product rather than retain them.

### Metabolic MAG-based models and community flux balance analysis reveal a minimal, stable, and highly functional community

GEMs were reconstructed for 13 MAGs selected for their predicted nutritional biosynthetic potential and metabolic functionality. The selection included *L. plantarum* 01 from kombucha, with predicted polysaccharide degradation capacity and SCFA biosynthetic potential; *L. plantarum* 02 from pozol, encoding over 40 CAZyme-annotated genes alongside high SCFA biosynthetic potential; *H. alvei* 01 and 02, both recovered from pulque, with notable predicted B12 and SCFA biosynthetic capacity; and *P. flexa* 01 and 02, which displayed the highest predicted SCFA biosynthetic potential across the entire dataset. *L. kefiranofaciens* 01 was selected for its high predicted SCFA production combined with B12 biosynthetic capacity in the absence of apparent B12 consumption; *L. kefiranofaciens* 02, while lacking distinctive metabolic features, was retained to enable intraspecific comparisons that may reveal functionally relevant strain-level variation within the same species. Two *L. kefiri* strains were also included, strain 01 showed the highest overall predicted probiotic potential, while strain 02 did not present distinguishing metabolic characteristics; nevertheless, both were retained for SynCom simulation to assess their respective contributions to community-level interactions. A similar rationale was applied to the yeast *B. bruxellensis*, where strain 01 displayed high predicted SCFA and B12 biosynthetic potential while strain 02 showed no outstanding metabolic profile, yet both were incorporated to capture the full spectrum of community interaction dynamics. Finally, *K. nataicola* from kombucha was included based on its predicted capacity to synthesize 1-heptadecene and to produce B12 without apparent consumption of this cofactor.

Supplemental Figure S4. provides genome-scale metabolic model summaries, including the number of reactions, metabolites, genes, exchange reactions, and model reconstruction statistics for each MAG. All models achieved optimal FBA solutions, with growth rates ranging from 25.4 to 117.0 mmol gDW⁻¹ h⁻¹ and model sizes ranging from 503 to 2,818 reactions and 454 to 1,853 metabolites (Table S1). Community metabolic simulations were performed using cooperative tradeoff FBA, which maximizes community-level growth while ensuring each member achieves at least 50% of its individual maximum growth rate.

Analysis of metabolic exchange fluxes across the 13-member consortium revealed a complex network of inter-organismal interactions dominated by amino acid and carbon source exchanges (Fig. 7A). *P. flexa* 01 and *H. alvei* 01 emerged as the primary donors of carbon sources, exporting ethanol, succinate, and galactose to multiple community members. Amino acid exchanges were widespread, with L-leucine, L-phenylalanine, and L-aspartate representing the most frequently exchanged metabolites across the community. Analysis of cofactor, nucleotide, and ion exchanges (Fig. 7B) revealed a prominent role for iron cycling, with siderophore-mediated iron transfer connecting multiple community members, suggesting cooperative iron acquisition strategies within the community.

**Figure 7.**
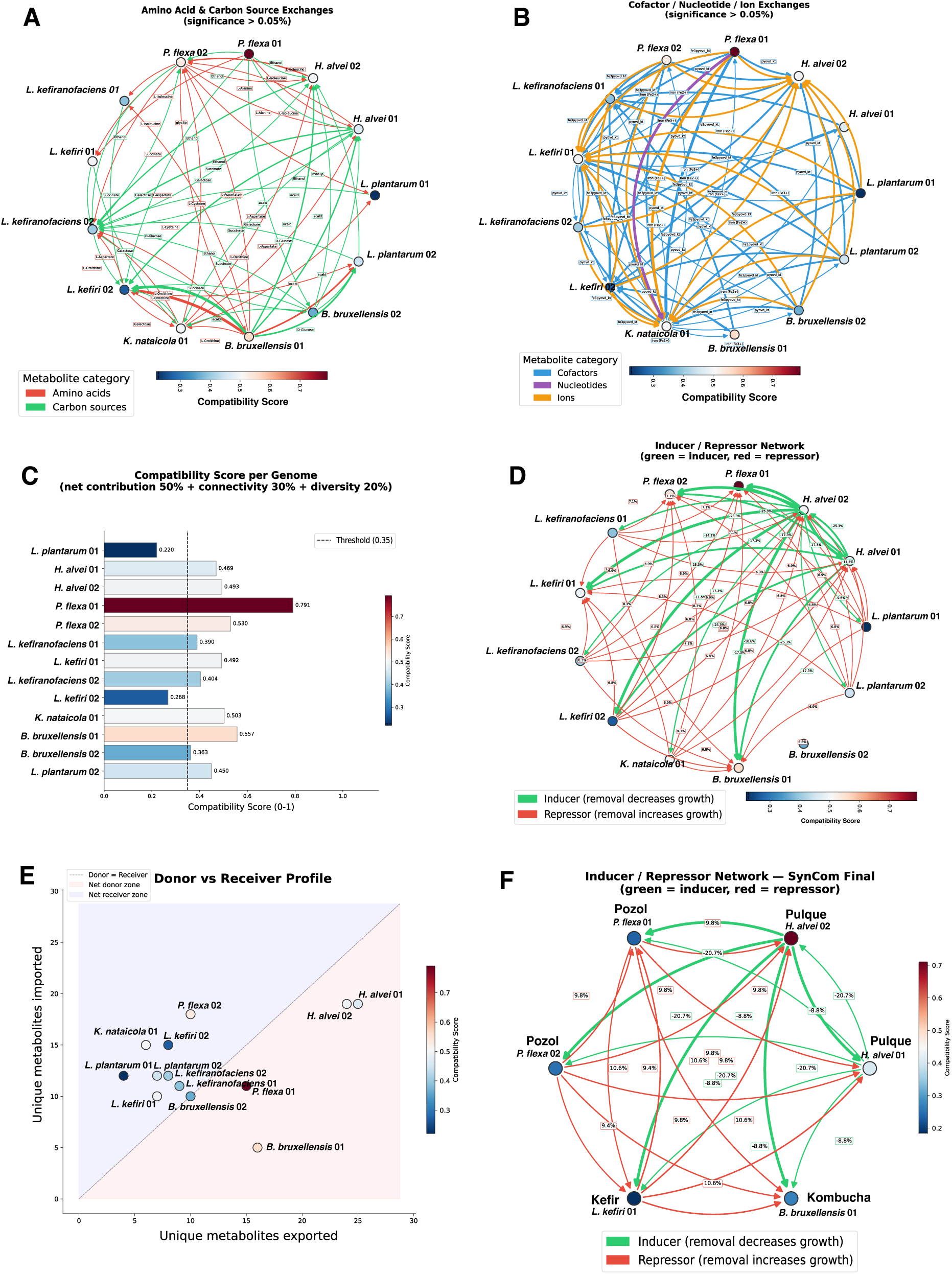
Metabolic interaction networks and community compatibility analysis of fermented beverage MAGs. (**A**) Amino acid and carbon source exchange network among community members (relative flux threshold > 0.05% of total organismal flux). Arrows indicate directional metabolite transfer between genomes; edge color denotes metabolite category (red = amino acids, green = carbon sources) and edge width reflects exchange flux magnitude. Node color indicates compatibility score. (**B**) Cofactor, nucleotide, and ion exchange network under the same significance threshold. Edge colors distinguish metabolite categories (blue = cofactors, orange = nucleotides, yellow = ions). (**C**) Compatibility score per genome, calculated as a weighted composite of net metabolic contribution (50%), metabolic connectivity (30%), and community diversity (20%). The dashed line indicates the selection threshold (0.35). (**D**) Inducer/repressor network for the full community based on leave-one-out analysis. Green arrows indicate induction (removal of source genome decreases growth of target by > 5%); red arrows indicate repression (removal increases growth by > 5%). Node color reflects compatibility score. (**E**) Donor vs. receiver metabolic profile. Each genome is positioned according to the number of unique metabolites exported (x axis) and imported (y axis). Genomes above the diagonal are net receivers; genomes below are net donors. (**F**) Inducer/repressor network of the final six-member SynCom shows the retained metabolic interaction topology. Edge labels indicate percent change in growth rate upon removal of the source genome. Green and red edges denote induction and repression, respectively.

*P. flexa* 01 achieved the highest compatibility score (0.791), followed by *B. bruxellensis* 01 (0.557) and *P. flexa* 02 (0.530), while *L. plantarum* 01 (0.220) and *L. kefiri* 02 (0.268) scored below the compatibility threshold of 0.35 (Fig. 7C). Leave-one-out community simulations were performed to identify induction and repression relationships among community members. Removal of each genome from the community and comparison of resulting growth rates against baseline revealed that *P. flexa* 01 functioned as the most critical inducer, with its removal causing growth rate reductions as low as −20.7% in multiple community members including *H. alvei* 01, *H. alvei* 02, *L. kefiri* 01, and *B. bruxellensis* 01 (Fig. 7D). Repressive interactions were also widespread, with several organisms showing increased growth rates upon removal of competitors, suggesting metabolic competition for shared resources.

Donor-receiver analysis (Fig. 7E) revealed that *B. bruxellensis* 01 was the most prolific metabolic donor, exporting a uniquely high number of metabolites relative to its imports, positioning it as a keystone metabolic contributor. In contrast, *H. alvei* 01 and *H. alvei* 02 showed high import profiles relative to exports, classifying them as net receivers within the community. Based on the integrated analysis of compatibility scores, metabolic exchange profiles, and inducer-repressor relationships, a minimal SynCom of six members was designed comprising *P. flexa* 01, *P. flexa* 02, *H. alvei* 01, *H. alvei* 02, *L. kefiri* 01, and *B. bruxellensis* 01. Community simulation of the SynCom confirmed stable cooperative growth, with *P. flexa* 01 maintaining its role as the primary inducer of community members (Fig. 7F). The SynCom retained the key metabolic exchange interactions observed in the full 13-member community while eliminating low compatibility and metabolically redundant members, resulting in a functionally coherent consortium predicted to support robust fermentation activity.

### Proposed workflow for the design of novel probiotic communities

In Figure 8A, we describe the complete workflow developed in this study for microbial mining of traditional fermented beverages to identify novel probiotic candidates. Shotgun sequencing FASTQ files were retrieved for each beverage, and reads underwent quality assessment, trimming, and assembly into high-quality MAGs, followed by completeness evaluation and taxonomic classification and metabolic modelling and prediction of the interaction network.

**Figure 8.**
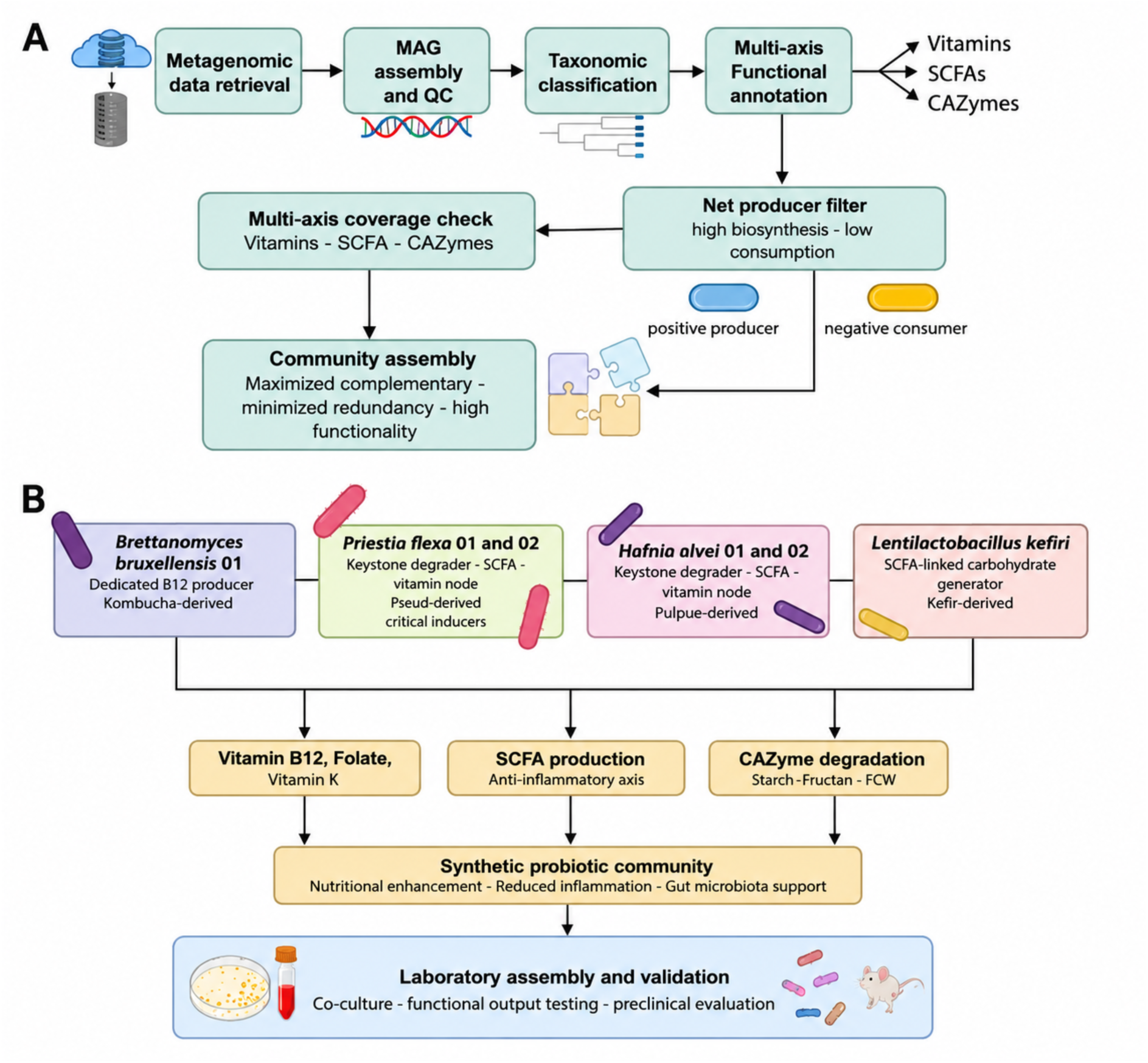
Methodology and rationale for the assembly of nutritionally enhanced, gut microbiota-modulating probiotic communities. (**A**) Bioinformatic workflow identifies genomic features guiding the assembly of highly functional microbial communities from shotgun metagenomic data of fermented beverages. Metagenome-assembled genomes (MAGs) were quality-filtered, taxonomically classified, and functionally annotated across three axes: vitamin biosynthesis, SCFA production, and CAZyme repertoires. Strains were selected based on net producer potential, maximizing biosynthetic capacity while minimizing metabolic redundancy. (**B**) Probiotic community assembled from the genomic data of the four fermented beverages analyzed in this study. The selected strains collectively cover vitamin biosynthesis (B12, Folate, and K), SCFA-driven anti-inflammatory activity, and degradation of complex carbohydrates including starch, fructans, and plant cell wall polysaccharides. The community is intended to enhance the nutritional value of a fiber-based functional beverage, promote intestinal acidification, and support gut microbiota homeostasis, with performance to be validated through co-culture experiments and preclinical evaluation.

We propose a core set of genomic criteria to assess probiotic potential. These comprise (I) vitamin biosynthetic capacity with low consumption of key micronutrients, particularly vitamin B12, for which microbial competition and analogue production can impair human absorption; (II) robust SCFA production, which contributes to pathogen reduction and intestinal anti-inflammatory activity; and (III) high CAZyme content, supporting degradation of complex carbohydrates and downstream nutrient release. A net production filter was applied to prioritize organisms with high biosynthetic and low consumption gene content across these axes.

Based on these criteria, we propose a six-member synthetic probiotic community characterized by high metabolic connectivity and strong functional interdependence (Figure 8B), selected to maximize functional complementarity while minimizing metabolic redundancy.

## Discussion

The exploration of fermented foods and beverages through shotgun metagenomics represents a novel and invaluable source of potential probiotic microorganisms, yet whose nutritional and health significance remains largely unexplored. Here, we analyzed available data from four independent studies that performed deep shotgun metagenomic sequencing on kombucha, kefir, pozol, and pulque. We found that despite diversity differences at the genus level, comparable numbers of MAGs were recovered from kombucha (24), pozol (25), and pulque (24), rendering these three beverages highly comparable at the genomic level regardless of library size. Notably, approximately 25% of MAGs from all three beverages could not be classified to any known species, suggesting they may represent previously undescribed bacteria. Undescribed microbial diversity has similarly been reported in other fermented food metagenomic surveys (Leech *et al.,* 2020). This finding reinforces the notion that traditional fermented beverages are unexplored reservoirs of novel microorganisms with potential probiotic properties. Consistent with this, taxonomic classification revealed a low degree of species overlap among beverages, highlighting the microbiological uniqueness of each fermentation process.

In pozol, a larger proportion of MAGs could be resolved to the species level compared to the other beverages. Additionally, we reconstructed the complete chloroplast genome of maize (*Zea mays*) from pozol. Although expected given the maize substrate, its successful reconstruction provides an independent validation of the metagenomic assembly pipeline and supports the quality of the recovered microbial genomes, a result not reported in the original pozol metagenomics study (López-Sánchez *et al.,* 2023), suggesting that our methodology achieves higher sensitivity for complete genome reconstruction.

Although the datasets used here are of high quality and derive from studies with clear and internally consistent conclusions, the unequal sample sizes across beverages (kefir n = 2, kombucha n = 23, pozol n = 5, and pulque n = 6) limit statistical comparison of cross-group diversity. Despite this constraint, the number of MAGs recovered was remarkably similar across three of the four beverages, which strengthens the comparability of the genome-resolved functional analyses that form the study’s core.

Kombucha was dominated by multiple species of *Komagataeibacter*, consistent with its established role as the primary acetic acid-and organic acid-producing genus, as well as the producer of the bacterial cellulose matrix that constitutes the characteristic SCOBY of this beverage (Subbiahdoss *et al.,* 2022; Su *et al.,* 2023). Kefir showed the most restricted composition, with only *Lentilactobacillus kefiri* and *Lactobacillus kefiranofaciens*, the principal fermentative agents of this beverage (González-Orozco *et al.,* 2022). *Zymomonas*, a genus broadly reported across fermented beverages, was also identified in our analysis, specifically in pulque. Interestingly, this same genus has been reported as a dominant member in other spontaneously fermented plant-derived beverages such as taberna, made from palm sap, which may reflect a shared substrate-driven selection associated with plant-derived sugars (Alcántara-Hernández *et al.,* 2010).

Several MAGs recovered in this study encode BGCs predicted by antiSMASH, a bioinformatics tool for identifying biosynthetic gene clusters, to produce compounds with documented biomedical activity. *K. rhaeticus* from kombucha emerged as the most biosynthetically remarkable organism, containing BGCs with high predicted similarity to clusters encoding both 1-heptadecene and 1-nonadecene. These are long-chain terminal alkenes with documented antifungal and broad-spectrum antimicrobial activity (Yoon *et al.,* 2024). In pozol, two organisms possessed BGCs with notable predicted biosynthetic potential. *Streptococcus infantarius* encoded a BGC with predicted similarity to thermophilin 1277, a lantibiotic-type bacteriocin with broad-spectrum antibacterial activity against lactic acid bacteria and food spoilage pathogens including *Clostridium* and *Bacillus cereus* (Kabuki *et al.,* 2007). The detection of alkene BGCs in *K. rhaeticus*, a bacterium primarily recognized for bacterial cellulose production, underscores the underappreciated secondary metabolic potential of this genus and the value of comprehensive functional annotation in metagenomic bioprospecting pipelines. A BGC with medium predicted similarity to the paeninodin cluster was also identified in a *Priestia flexa* 02-MAG from pozol. Paeninodin is a ribosomally synthesized lasso peptide first described in *Paenibacillus dendritiformis*, notable for using a unique kinase-mediated phosphorylation tailoring step and reported to exhibit antimicrobial activity (Zhu *et al.,* 2016).

*Hafnia alvei* 02 from pulque encodes a BGC with high predicted similarity to desferrioxamine E, a hydroxamate siderophore with exceptionally high Fe³⁺ binding affinity. Siderophore production by *H. alvei* is mechanistically relevant in the context of its broader metabolic profile. Cobalamin biosynthesis requires iron-dependent enzymatic steps and cobalt chelation through a corrin ring structurally related to porphyrin precursors shared with heme and siroheme (Balabanova *et al.,* 2021), suggesting that this organism can mobilize environmental iron through siderophore-mediated chelation may directly sustain its predicted B12 biosynthetic potential.

Metabolic reconstruction through CAZyme profiling revealed substrate-driven functional specialization across beverages. In pozol, starch-degrading enzymes were prominently represented, as expected given nixtamalized maize as the fermentation substrate (Díaz-Ruiz *et al.,* 2003), suggesting these bacteria could also support dietary starch digestion upon intestinal colonization. In pulque, fructan-and inulin-degrading enzyme families were enriched, reflecting the fructooligosaccharide-rich composition of agave sap (Escalante *et al.,* 2016). Of particular note, two MAGs assigned to *P. flexa* 01 and 02 from pozol combined high CAZyme biosynthetic capacity with predicted efficient production of SCFAs, positioning this species as a functionally relevant probiotic candidate.

Propionate, a key SCFA, was identified as a differentially produced metabolite in kombucha bacteria. SCFAs produced through fermentation of inulin, pectin, resistant starch, and fructooligosaccharides by gut bacteria such as *Bacteroides* spp., *Faecalibacterium prausnitzii*, and *Akkermansia muciniphila* are known to nourish colonocytes, maintain intestinal barrier integrity, regulate immune responses, particularly regulatory T cells, and modulate appetite hormones GLP-1 and PYY (Canfora *et al.,* 2019). These findings suggest that kombucha-associated bacteria capable of producing propionate may confer meaningful metabolic benefits upon intestinal colonization (Chambers *et al.,* 2015). Also, *P. flexa* 01/02 and *H. alvei* 01 from pozol and pulque also appear to be strong producers of these molecules, suggesting they may exert similar effects on intestinal barrier protection and beneficial outcomes in obesity (Legrand *et al*., 2020).

Beyond carbohydrate metabolism, bacterial vitamin biosynthesis was detected across all beverages. Consistent with prior reports, kombucha bacteria showed strong B-vitamin biosynthetic potential (Jayabalan *et al.,* 2014). Strikingly, both the pozol and pulque microbiota exhibited notable genetic capacity for folate (B9) and vitamin K biosynthesis. While folate biosynthetic potential in pulque has been previously reported at the functional gene level (Chacón-Vargas *et al.,* 2020), and general vitamin biosynthetic capacity has been described in pozol, the specific capacity for vitamin K biosynthesis in these beverages remains largely unexplored.

The comparative analysis across all MAGs identified specific microbial candidates combining substrate degradation capacity with biosynthesis of health-relevant compounds with properties that would be more efficiently delivered as isolated strains rather than as part of complex artisanal consortia. *P. flexa* 01 and 02 (pozol) and *H. alvei* 01 and 02 (pulque) stood out as top candidates; the latter encodes biosynthetic pathways for multiple B vitamins and SCFAs (Petraro *et al.,* 2024), and clinical supplementation with the strain *H. alvei* HA4597 was associated with greater weight loss and reduced hunger compared to a placebo (Déchelotte *et al.,* 2021). In kombucha, the yeast *B. bruxellensis* emerged as a standout candidate, combining high complex carbohydrate degradation capacity, elevated B12 biosynthetic potential with low internal B12 consumption, and SCFA production, a functional profile consistent with its confirmed presence as a core member of the kombucha consortium (Liao *et al.,* 2024). The probiotic relevance of pulque-derived *Lactiplantibacillus plantarum* strains is further supported by experimental evidence. A strain isolated from pulque produced in Morelos, Mexico was recently shown to resist acidic pH and bile salts, reduce cholesterol, display broad-spectrum antimicrobial activity against both Gram-positive and Gram-negative pathogens, and confer *in vivo* protection against *Salmonella* infection in a murine model; genome mining also revealed the presence of plantaricin EF and other bacteriocin-encoding genes (Giles-Gómez *et al.,* 2024). Compatibility scoring within the SynCom framework, however, did not favor *L. plantarum*; instead, *Lentilactobacillus kefiri* emerged as a highly central member of the designed consortium. *L. kefiri*, one of the most characteristic bacteria found in kefir grains, has demonstrated resistance to bile salts and antimicrobial activity against pathogens including *Bacillus cereus* and *Staphylococcus aureus*. It also possesses the capacity to attenuate proinflammatory responses in intestinal epithelial cells through a high IL-10/IL-12 ratio, supporting its dual role as an antimicrobial and immunomodulatory probiotic agent (Carasi *et al.,* 2014; Carasi *et al.,* 2015).

The metabolic exchange networks of these strains revealed a rich landscape of intermicrobial micronutrient sharing, encompassing amino acids, carbon sources, cofactors, nucleotides, and ions, underscoring the functional interdependence of the assembled community. Notably, cofactor and ion exchange emerged as particularly prominent interaction categories, suggesting that micronutrient cross-feeding, rather than competition for shared substrates, is a primary driver of community cohesion in this SynCom. A striking feature of the final six-member community is the inclusion of two conspecific strain pairs, *Hafnia alvei* 01/02 and *Priestia flexa* 01/02, both of which exceeded the compatibility threshold and were retained through leave-one-out inducer-repressor profiling. This outcome suggests that intraspecific strain diversity contributes positively to community robustness, likely through complementary metabolic niches that minimize direct competition while maintaining functional redundancy in key roles such as B12 biosynthesis and CAZyme-mediated carbohydrate degradation. From a bioprospecting perspective, this finding carries practical implications, when isolating microbial candidates from traditional fermented beverages, recovering multiple strains of the same species may be as strategically valuable as expanding taxonomic breadth, particularly for species with demonstrated probiotic potential such as *Hafnia alvei* and *Priestia flexa*. This pattern aligns with observations in other microbial ecosystems, where within-species genetic diversity expands as community diversity increases, driven by additional metabolic niches that allow multiple strains to coexist without full functional overlap (Madi *et al.,* 2023). Such strain-level variation carries ecological consequences not predictable from species identity alone, supporting the value of capturing intraspecific diversity when assembling microbial consortia. The retention of *L. kefiri* 01 and *B. bruxellensis* 01 as non-redundant members further highlights the value of cross-kingdom diversity, combining bacterial and yeast metabolisms to achieve a stable, highly connected consortium that sustains cooperative growth across a broad metabolic repertoire.

A mechanistically compelling interaction emerging from the metabolic exchange network is the functional axis between *P. flexa* and *H. alvei*. Members of the *Priestia* genus have well-characterized siderophore biosynthesis gene clusters, capable of mobilizing iron within microbial communities (Zhu *et al.,* 2025). Consistently, *P. flexa* emerged as a net ion donor in our community flux balance simulations. This is relevant because cobalamin biosynthesis shares the common precursor precorrin-2 with siroheme and heme, both of which involve iron-dependent enzymatic steps and a centrally chelated metal ion (Balabanova *et al.,* 2021). Accordingly, *H. alvei* 01, the predicted B12 producer in the consortium, was a net ion acceptor, suggesting that iron supplied by *P. flexa* directly sustains its cobalamin biosynthetic capacity. Siderophore-mediated iron cross-feeding has been documented as a mutualistic interaction in the gut microbiome, where iron donor species actively enhance the metabolic performance of iron-dependent partners (Culp *et al.,* 2023). We propose that this *P. flexa*-*H. alvei* axis constitutes a keystone interaction within the SynCom, linking iron homeostasis to B12 availability and providing a mechanistic rationale for retaining both species, and their conspecific strain pairs, in the final community design.

Although kombucha and kefir have been widely promoted as highly beneficial beverages, this comparative bioinformatic analysis revealed that traditional spontaneously fermented beverages may contain microorganisms with greater probiotic potential, while also displaying high microbial diversity. Based on these findings, we propose criteria for the systematic exploitation of shotgun metagenomic data from underexplored fermented foods and beverages. These include MAG reconstruction with an inclusive scope that encompasses not only bacteria but also yeasts and fungi; the functional analysis of recovered genomes through metabolic reconstruction to identify competitive and inductive interactions; and the *in silico* design of SynComs through community-level metabolic modeling. Ultimately, the health-promoting potential of candidate strains and synthetic communities identified through this framework will require validation through controlled in vitro studies, animal models where appropriate, and well-designed human clinical trials.

## Author Contributions

JMVE conceived the study. JMVE, AT-G, AAR-H, and JTO-P designed the preliminary versions of the project. AT-G collected the initial metagenomics public data and performed the analysis including taxonomic classification, assembly analyses, BUSCO, and functional annotation. AAR-H developed the baseline computational scripts, contributed to data generation, and designed and refined all figures and visualizations. JTO-P and COV-R performed the short-chain amino acid analyses. JMVE, AT-G, and AAR-H wrote the initial draft of the manuscript. JMVE and AAR-H led the text revisions, with significant writing contributions from AT-G and JTO-P. IPC and AGA reviewed the manuscript for quality and clarity. All authors read and approved the final manuscript.

## Acknowledgments

The authors would like to thank the Institute for Obesity Research at the Monterrey Institute of Technology for providing the institutional support and resources to carry out this study. We are also deeply grateful to Dr. Nelly Selem Mojica from the Center for Mathematical Sciences at the National Autonomous University of Mexico for granting access to their high-performance computers and servers, essential for our data processing and bioinformatic analyses. AAR-H acknowledges SECIHTI for the doctoral fellowship granted during the development of this research. Finally, we thank John Kohl for his careful and insightful review of the manuscript.

## Data Availability Statement

This studýs assembled and analyzed MAGs have been deposited in Zenodo under accession https://doi.org/10.5281/zenodo.21210858. Bioinformatics pipelines and analysis scripts are available at https://github.com/alexandratre1/Bebidas-Fermentadas/blob/main/README.md.

