## Supplementary_information for "Genome-resolved metagenomics of traditional fermented beverages reveals biosynthetic diversity and informs the rational *in silico* design of probiotic synthetic communities"

**Table of contents** **1**

**Supplementary Figures** **2**

Figure S1. Geographic distribution of the fermented beverages included in this study. 2

Fig S2. Distribution of vitamin biosynthesis and uptake routes per taxon across fermented beverages. 3

Fig S3 Mean total counts of cobalamin metabolism gene routes across taxa and beverages. 4

Figure S4. Distribution of enriched genes based on each functional category. 5

Fig. S5. Metabolic model summaries of representative metagenome-assembled genomes (MAGs). 6

**Supplementary Tables** **7**

**Supplementary Figures**

S1. Geographic distribution of fermented beverages

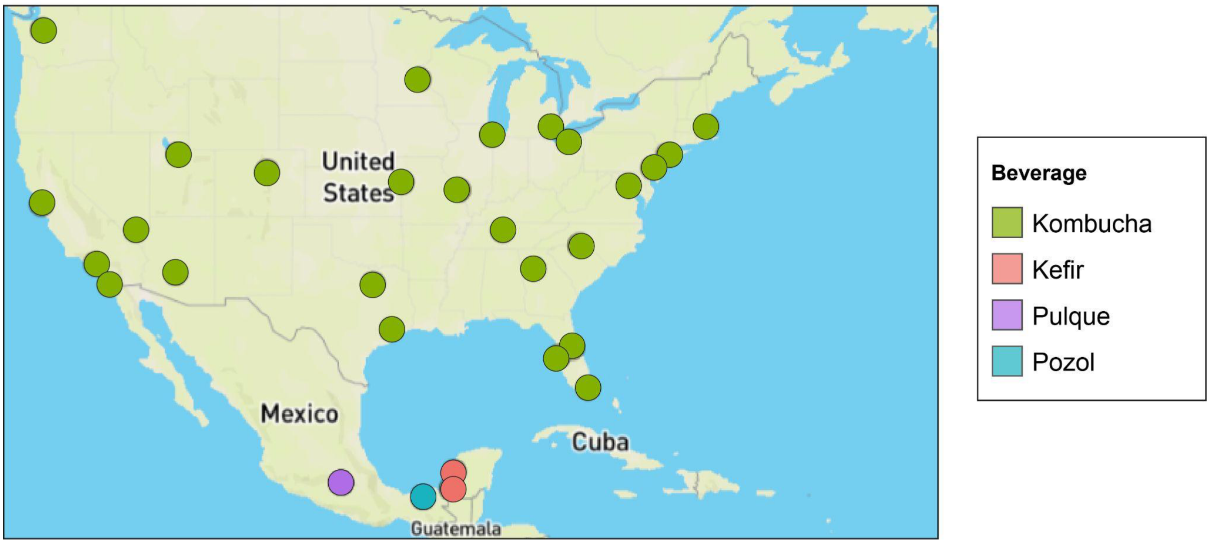


**Figure S1. Geographic distribution of the fermented beverages included in this study.**
Sampling locations corresponding to the four fermented beverage types analyzed are shown across North America. Colored markers indicate the origin of metagenomic datasets obtained from kombucha (green), kefir (pink), pulque (purple), and pozol (cyan). Kombucha samples were retrieved from a broad range of locations throughout the United States, whereas kefir, pulque, and pozol samples were primarily obtained from Mexico and Guatemala, reflecting the traditional geographic distribution of these fermented beverages. This dataset encompasses diverse ecological and cultural origins, providing a representative framework for comparative metagenomic analyses of microbial community composition and functional potential.

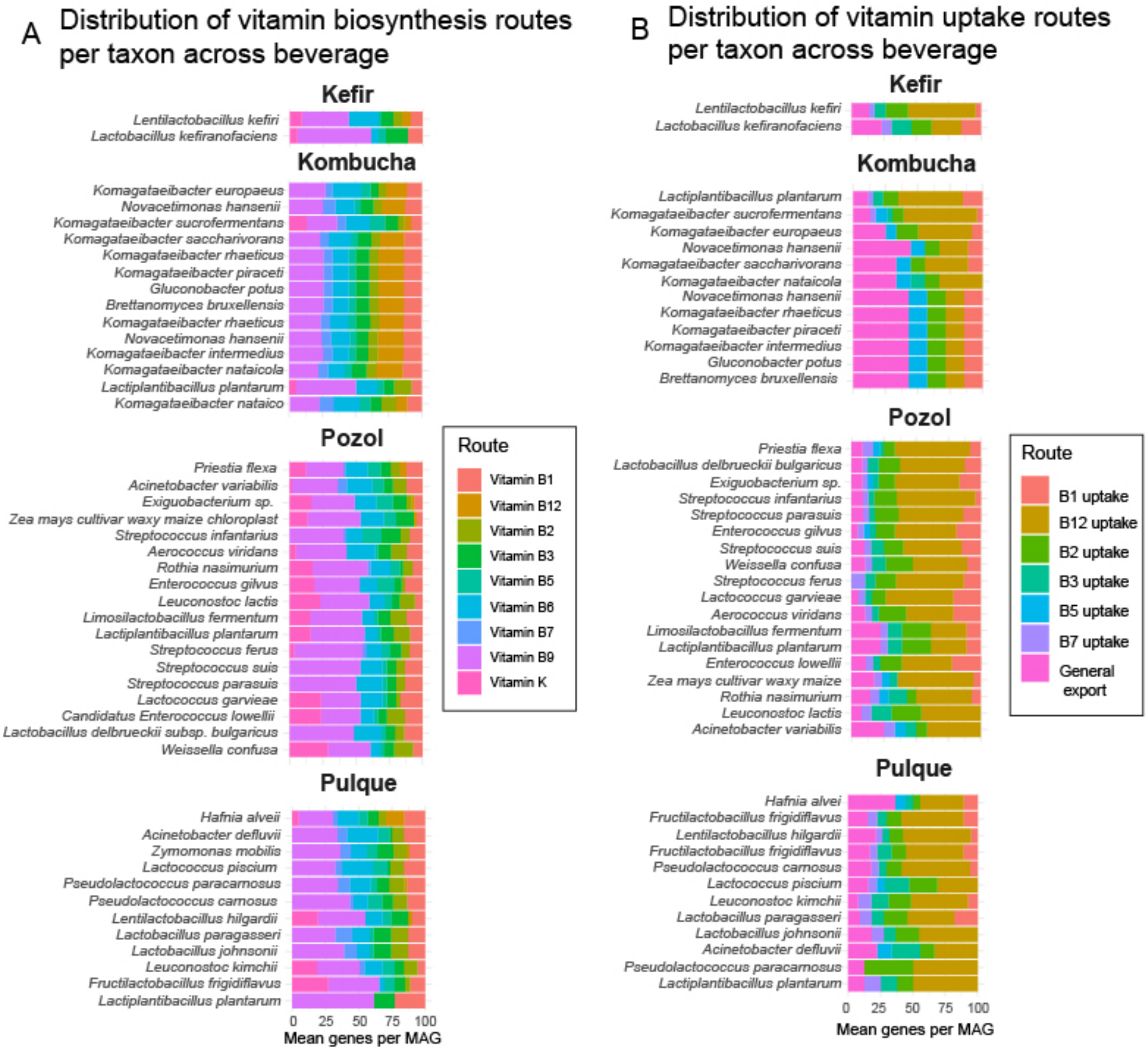


**Fig S2. Distribution of vitamin biosynthesis and uptake routes per taxon across fermented beverages.**

The figure presents paired 100% stacked horizontal bar charts illustrating the relative proportion of specific vitamin-related pathways within the overall genetic repertoire for vitamin metabolism in dominant taxa, grouped by beverage of origin (Kefir, Kombucha, Pozol, and Pulque). (**A**) Distribution of vitamin biosynthesis routes. The chart details the relative genomic contribution of nine distinct vitamin biosynthesis pathways (Vitamins B1, B12, B2, B3, B5, B6, B7, B9, and K), each denoted by a specific color according to the legend. The profiles reveal that Vitamin B9 biosynthesis (light purple) accounts for a major proportion of the synthesis repertoire in most taxa across Kefir, Pozol, and Pulque. In contrast, Kombucha-associated taxa display a distinct profile with a notable proportion dedicated to Vitamin B12 biosynthesis. (**B**) Distribution of vitamin uptake and export routes. The chart shows the relative proportion of uptake pathways for specific vitamins (B1, B12, B2, B3, B5, B7) and general export. Unlike biosynthesis, the distribution of uptake routes shows that B12 uptake constitutes a substantial and ubiquitous proportion of the transport genetic repertoire across almost all evaluated taxa, regardless of their beverage of origin. The X-axis in both panels is scaled from 0 to 100, representing the relative percentage of each route within the total identified vitamin metabolism genes per metagenome-assembled genome (MAG).

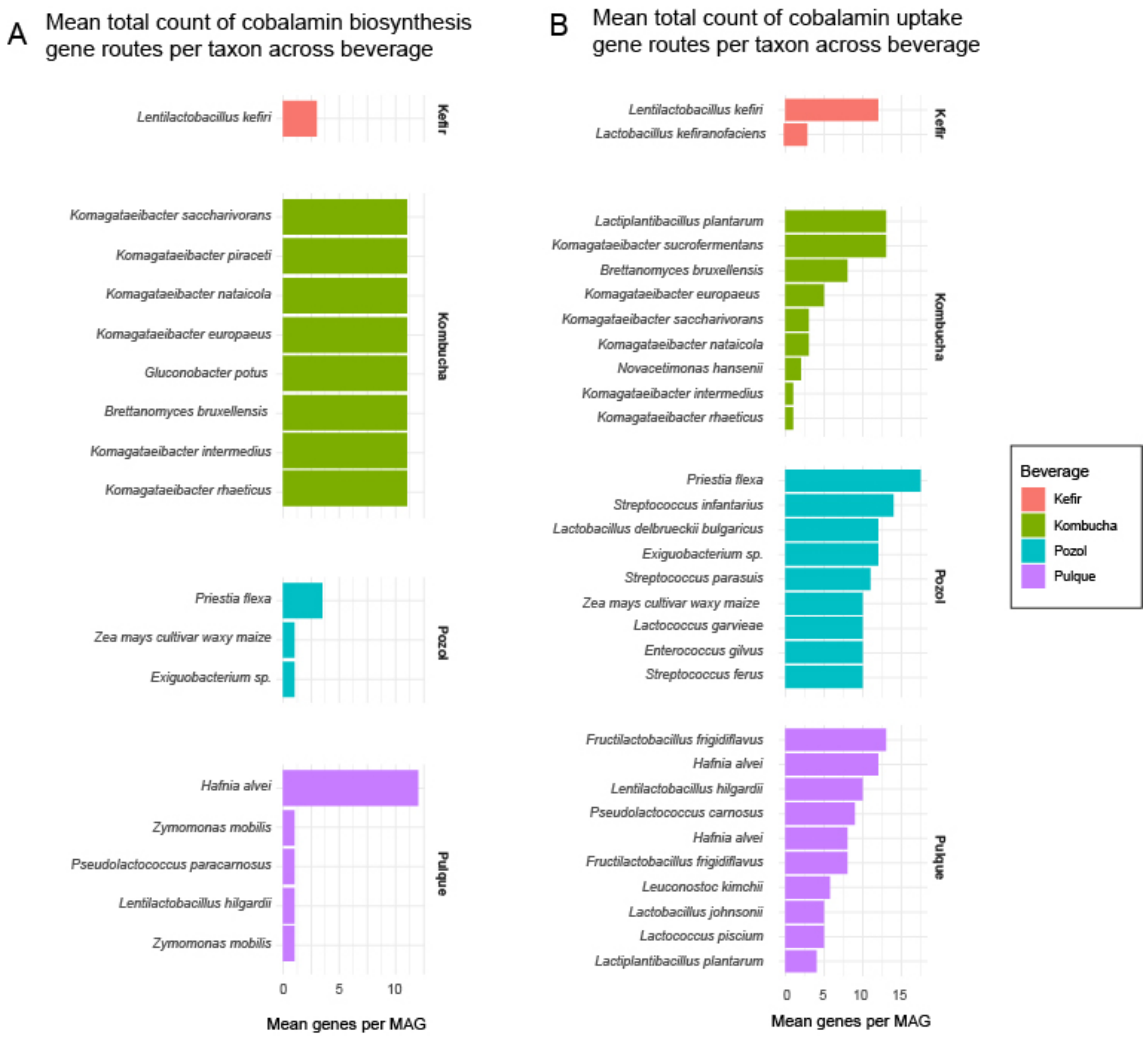


**Fig S3 Mean total counts of cobalamin metabolism gene routes across taxa and beverages.**
The figure displays paired horizontal bar charts for two aspects of cobalamin (vitamin B12) metabolism across relevant taxons, stratified by beverage of origin, colored according to the legend. (**A**) Chart detailing the mean total count of genes involved in cobalamin biosynthesis routes for each taxon. High biosynthesis potential is concentrated among taxa associated with Kombucha, including *Komagataeibacter* species, *Gluconobacter potus*, and *Brettanomyces bruxellensis*. Conversely, most taxa in other beverages have low biosynthesis counts, with the exception of *Hafnia alvei* strains from Pulque which show counts of 8-12 genes. (**B**) Chart detailing the mean total count of genes for cobalamin uptake routes for the same taxa. High uptake potential is found in a diverse array of taxa from all beverage types, with many MAGs exhibiting counts of 10-14 genes. This includes widespread capacity in Lactic Acid Bacteria from Kefir and Pozol, and various species in Kombucha and Pulque. The overall pattern highlights that while cobalamin biosynthesis capacity is taxonomically and beverage-specific, cobalamin uptake is a more ubiquitous feature across the diverse microbiota of these fermented beverages. Taxon names are listed individually on the Y-axis. The X-axis for both charts represents the mean number of genes per metagenome-assembled genome (MAG).

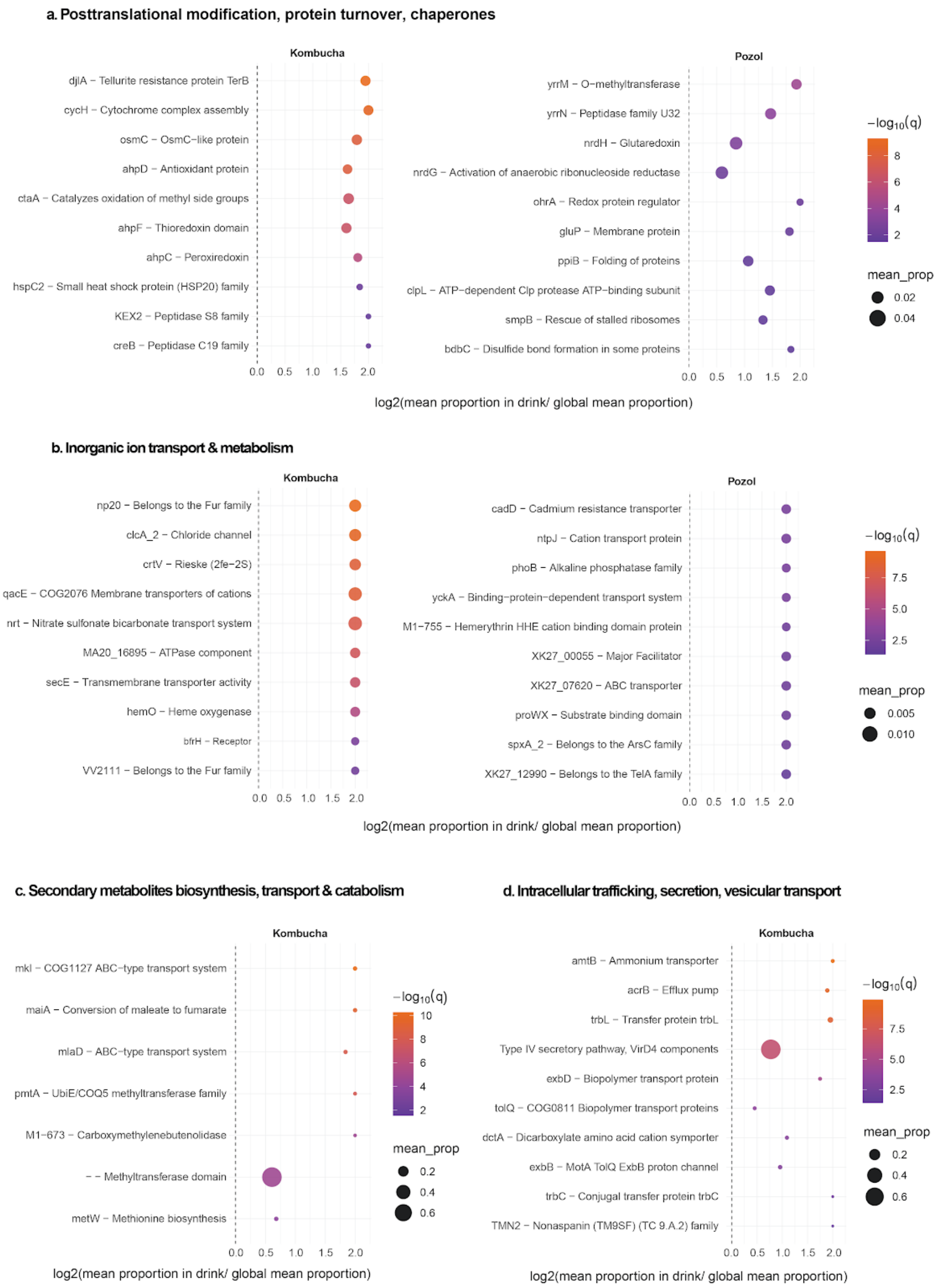


**Figure S4. Distribution of enriched genes based on each functional category.**
Taking into account Figure 4, this supplementary image highlights certain functional categories of vital relevance to the microorganisms present in the beverage and to the people who consume it. (a) The category of post-translational modification, protein turnover, and chaperones is shown, where kombucha exhibits greater enrichment in genes related to oxidation, the response to heat stress, and peptide consumption; while pozol focuses on protein formation, the response to oxidation, and membrane preservation. (b) In the category of inorganic ion transport and metabolism, kombucha is again observed, now with overexpression of proteins for iron and oxygen recovery, as well as ATP production; pozol is also present, but this time as proteins of resistance to cadmium and toxic ions in general. (c) In the enrichment of secondary metabolites, transport, and catabolism, kombucha is the only beverage with overall enrichment, demonstrating primarily genes for methylation and substrate conversion. (d) Kombucha is again the only beverage with notable values in intracellular trafficking,

secretion, and vesicular transport, particularly in genes that transport ammonia, biopolymers, and cations.

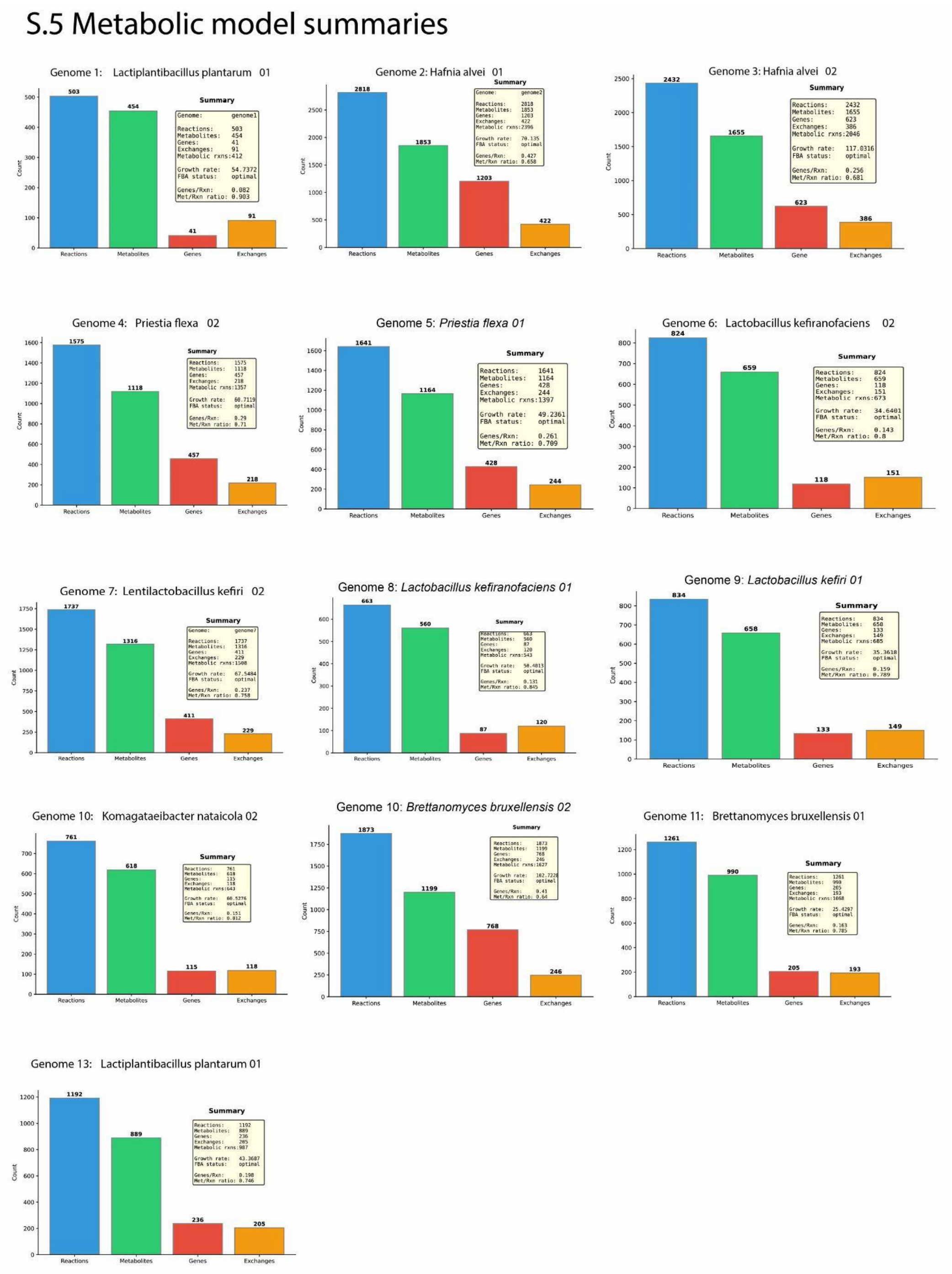


**Fig. S5. Metabolic model summaries of representative metagenome-assembled genomes (MAGs).** The figure presents individual bar charts summarizing the reconstructed genome-scale metabolic models for representative MAGs from fermented beverages. Each panel corresponds to a single genome and displays the total number of reactions, metabolites, genes, and exchange reactions included in the metabolic reconstruction. An inset summary accompanies each model, reporting additional reconstruction metrics, including the total number of metabolic reactions, predicted growth rate, flux balance analysis (FBA) status, and gene-to-reaction and metabolite-to-reaction ratios. The reconstructed models encompass bacterial and yeast genomes representative of Kefir, Kombucha, Pulque, and Pozol microbiomes, including *Lactiplantibacillus plantarum, Hafnia alvei, Priestia flexa, Lactobacillus kefiranofaciens, Lentilactobacillus kefiri, Komagataeibacter nataicola,* and *Brettanomyces bruxellensis*. Overall, the figure provides a comparative overview of the size and composition of each genome-scale metabolic model, highlighting differences in metabolic network complexity among reconstructed genomes.

**Supplementary Tables**

1. **Supplementary Table 1**. Metadata and sequencing statistics of fermented beverage metagenomic samples.
2. **Supplementary Table 2.** Geographic location of all beverage samples.
3. **Supplementary Table 3.** Taxonomic classification of metagenomes for all beverages
4. **Supplementary Table 4.** BUSCO completeness of recovered MAGs.
5. **Supplementary Table 5.** Taxonomic classification of the high-quality recovered genomes for all beverages.
6. **Supplementary Table 6.** Short fatty acids contribution of each genome.
7. **Supplementary Table 7.** Cobalamin contribution of each genome.
8. **Supplementary Table 8**. Metabolite model analysis.
9. **Supplementary Table 9.** Metabolic model summaries.
